# Collective synchrony of mating signals modulated by ecological cues and social signals in bioluminescent sea fireflies

**DOI:** 10.1101/2023.06.16.545275

**Authors:** Nicholai M. Hensley, Trevor J. Rivers, Gretchen A. Gerrish, Raj Saha, Todd H. Oakley

## Abstract

Individuals often employ simple rules that can emergently synchronise behaviour. Some collective behaviours are intuitively beneficial, but others like mate signalling in leks occur across taxa despite theoretical individual costs. Whether disparate instances of synchronous signalling are similarly organised is unknown, largely due to challenges observing many individuals simultaneously. Recording field collectives and *ex situ* playback experiments, we describe principles of synchronous bioluminescent signals produced by marine ostracods (Crustacea; Luxorina) that seem behaviorally convergent with terrestrial fireflies, and with whom they last shared a common ancestor over 500 mya. Like synchronous fireflies, groups of signalling males use visual cues (intensity and duration of light) to decide when to signal. Individual ostracods also modulate their signal based on the distance to nearest neighbours. During peak darkness, luminescent “waves” of synchronous displays emerge and ripple across the sea floor every ∼60 seconds, but such periodicity decays within and between nights after the full moon. Our data reveal these bioluminescent aggregations are sensitive to both ecological and social light sources. Because the function of collective signals is difficult to dissect, evolutionary convergence, like in the synchronous visual displays of diverse arthropods, provides natural replicates to understand the generalities that produce emergent group behaviour.

## Introduction

Animal collectives perform some of the most striking examples of behaviour observed [1]. Surprisingly, self-organised aggregations can be the emergent product of individuals following simple behavioural rules [2], as opposed to those that require complicated cognition or decision-making to form. Within these groups, individuals may use these heuristics at local spatial and temporal scales, influencing their own behaviour. For example, individual birds within a flock simply track a number of neighbours (i.e. topological [3]) or the distance to nearest neighbours (i.e. metric [4]) in order to coordinate individual flight heading and speed during murmurations. The form and strength of these local mechanisms produces variation in the collective.

Beyond how local interactions scale up, collectives may also be responsive to the environment [5]. Understanding how environmental variables shape variation in animal groups is a major goal in understanding collective dynamics [6,7]. For example, calling cicadas are sensitive to environmental light levels, where spatial synchrony within a forest increases when sound amplitude peaks as individuals increase their calling rate during brighter illumination [8]. However, tracking individuals within groups across environments to tackle such questions is only beginning to gain methodological traction. By understanding the balance between which local rules individuals use [9] and environmental effects, we can model variable patterns of collective action like decision making [10], swarming [11], or synchronisation [12,13].

Synchrony is generally considered the proximity in time of animal behaviours [2]. In animal communication, synchronous mate signalling represents an evolutionary paradox: in a classic scenario with males signalling for the attention of females, many males may aggregate and signal closely in space (called “leks”) and/or time (“sprees”) [14]. Although this cacophony of signals should be able to recruit more females from farther away (i.e. beacon effect hypothesis [15]), increasing the density of signals increases costs (i.e. more competition, hinders choice) [16]. Despite these, synchronous displays are found in diverse taxa and modalities [17]. Just across arthropods, synchronised behaviours are taxonomically scattered and probably arose multiple times from within groups without synchronous displays, suggesting convergent evolutionary events [18,19]. Outside the defensive visual displays of lepidopteran caterpillars [20] and vibrations of treehoppers [21], synchronised mating signals are known from choruses of singing orthopterans [22,23] and cicadas [8], the waving claws of fiddler crabs [24,25], and most famously, the glowing, flash bursts of synchronous fireflies found in Southeast Asia and the Americas [26,27]. These fireflies differ from their non-synchronous relatives, and as in other collective behaviours, seem to use local rules such as responding to the timing [28,29] of visual neighbours [27], which produces emergent synchrony. Although not conclusive, convergence may point towards similar evolutionary pressures (i.e. selection for competitive ability, or group-level adaptive value). Here, we add to the collection of synchronous visual displays by documenting the collective bioluminescent behaviours of shallow-water ostracods that show extreme synchrony [30]. By characterising this charismatic and independently evolved behaviour, we propose that luxorine sea fireflies [31] are an excellent system to study the causes and consequences of collective signalling.

Ostracods are millimetre-sized crustaceans found in fresh and marine waters, and many species possess a number of specialised adaptations such as thermophilia [32], desiccation-resistant resting eggs [33], or bioluminescence [34]. Males of most known species within the subtribe Luxorina, almost exclusively found in the reefs and seagrass beds of the Caribbean Sea [35,36], create distinct patterns of bioluminescence to attract females by secreting discrete packets of protein and substrate outside their body from a specialised glandular organ and into the water [37–39]. Females use these species-specific displays to orient and swim towards males [40,41], never flashing back. Besides visible displays, males also readily and rapidly switch behavioural tactics based on their proximity to the displays of other males, as well as the density of competitors: in the species *Photeros annecohenae [42,43]* males will either begin to display in close proximity and time to another male (termed “entrainment”) or sneak on another display by following along without producing light [44]. Signalling activity of a population is coincident with levels of available darkness during the lunar cycle [45], indicating that both social and ecological factors are important in modulating individual signalling behaviour..

To investigate the role of ecological and social factors in shaping collective synchronisation, we performed *in situ* observations and *ex situ* experiments using a species of ostracods within the same genus *Photeros* [31,43]. Currently known by its field designation “EGD” (hereafter), this species lives in seagrass beds off the coast of Panama, and upon first documentation, seemed extremely susceptible to stimulation with external stimuli, like the signals of other males, flashlights, lightning, etc. Anecdotally, many hundreds of animals would initiate their displays within seconds of one another, causing a cascade of light to ripple across the sea floor as new males began their displays before previous ones faded (Fig. 1B). These waves could extend for at least 10 metres, a collective feat for individuals less than 2 mm in size. Although we lacked the ability to accurately measure their spatial extent in the field, we sought to quantify these initial observations. Our data reveal that EGD males are sensitive to the level and timing of external light cues, resulting in observed waves of collective behaviour. Through field recordings, we also observe that ecological opportunity influences variation in these collectives. By quantifying and comparing these rules with those of other synchronous taxa like fireflies, we can begin to generalise on the shared and unique principles underlying emergent behaviours like collective signalling.

**Figure 1.**
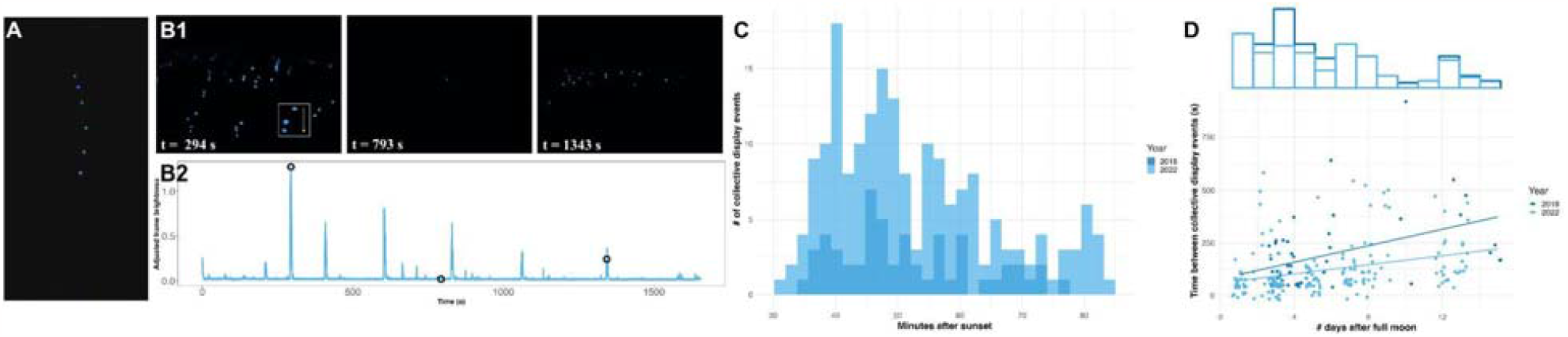
EGD displays are collectively periodic throughout the nights of the lunar cycle. (A) A single EGD display train filmed on a Canon ME20 with a Canon EF 16-35mm, f/2.8L III USM, inside a Gates underwater housing. Image by Presley Adamson and Christy Chamberlain, Monterey Bay Aquarium. (Bi) Video frames (false colour) of EGD signals captured using a WATEC 910HX camera in a custom underwater housing for ∼30 min. during a single night of observation, showing bouts of collective behaviour in the wild. From left to right: peak, during an interwave interval, and a smaller, more distant wave. Inset in (B) highlights the display of a single male with an added arrow to indicate the inferred swimming direction. (Bii) Total brightness across all pixels per frame over time, adjusted with a gamma correction per frame for changes in dynamic sensitivity. Black circles correspond to time points in (B1) from left to right. Available nighttime darkness limits the signalling niche within (C) a night and (D) a lunar cycle. (C) The number of collective behavioural events (y-axis) across field seasons decreases later into the night (x-axis), as nautical twilight ends. (D) Time between collective events (y-axis) increases days out from the full moon (x-axis), shown for two field seasons (separate regression lines) and with a marginal histogram depicting the total number of collective events per night. Year of field season indicated by shade: 2018 (dark), 2022 (light).

## Results

### EGD displays are distinct from other Luxorina species

Like other ostracod species in the subtribe Luxornia, individual males of the species EGD produce spatiotemporally complex signals (Fig. 1A). EGD swims downwards while producing a series of 4 - 8 discrete pulses that last 2 - 5 sec each in a line (Table 1; Fig. 2A). Because the interpulse intervals are ∼1.5 sec long, multiple pulses are visible simultaneously as a male secretes them and continues to swim towards the ocean floor. Compared to other *Photeros* species, EGD has intermediate pulse durations and interpulse intervals, with durations that are longer than ‘flashing’ species like *P. morini* [37,46] but much shorter than others [43]. They are also spatially short, with 4 - 8 cm between pulses, and the overall display length is very compact, covering only ∼16 cm compared to other *Photeros* that range from 20 – 180 cm (Fig. S1). As with other Luxorine species, the relative intensity of subsequent pulses decreases throughout a display, but our low sample sizes should be interpreted as a preliminary estimate.

**Table 1.**
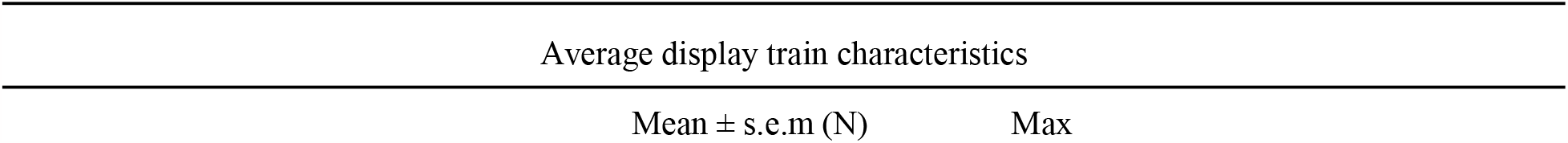
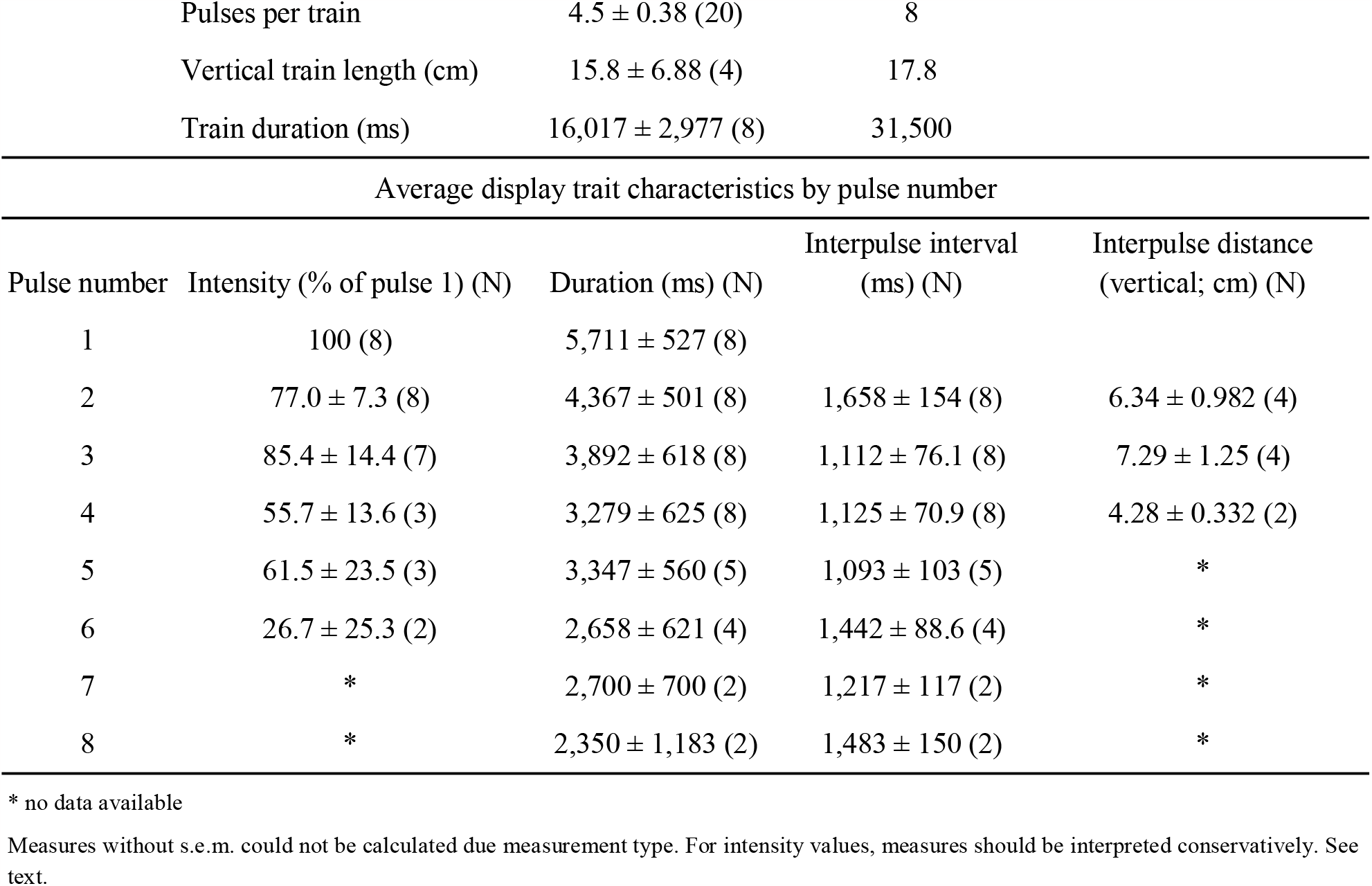
EGD display characteristics from both stereoscopic and single camera recordings *in situ*. Pulse duration and interpulse interval are in ms. Distances are in cm. Intensity is relative, normalised to the first pulse within a display train. Interpulse intervals and distances begin between pulse 1 and 2. Replicates are in parentheses, *N*.

**Figure 2.**
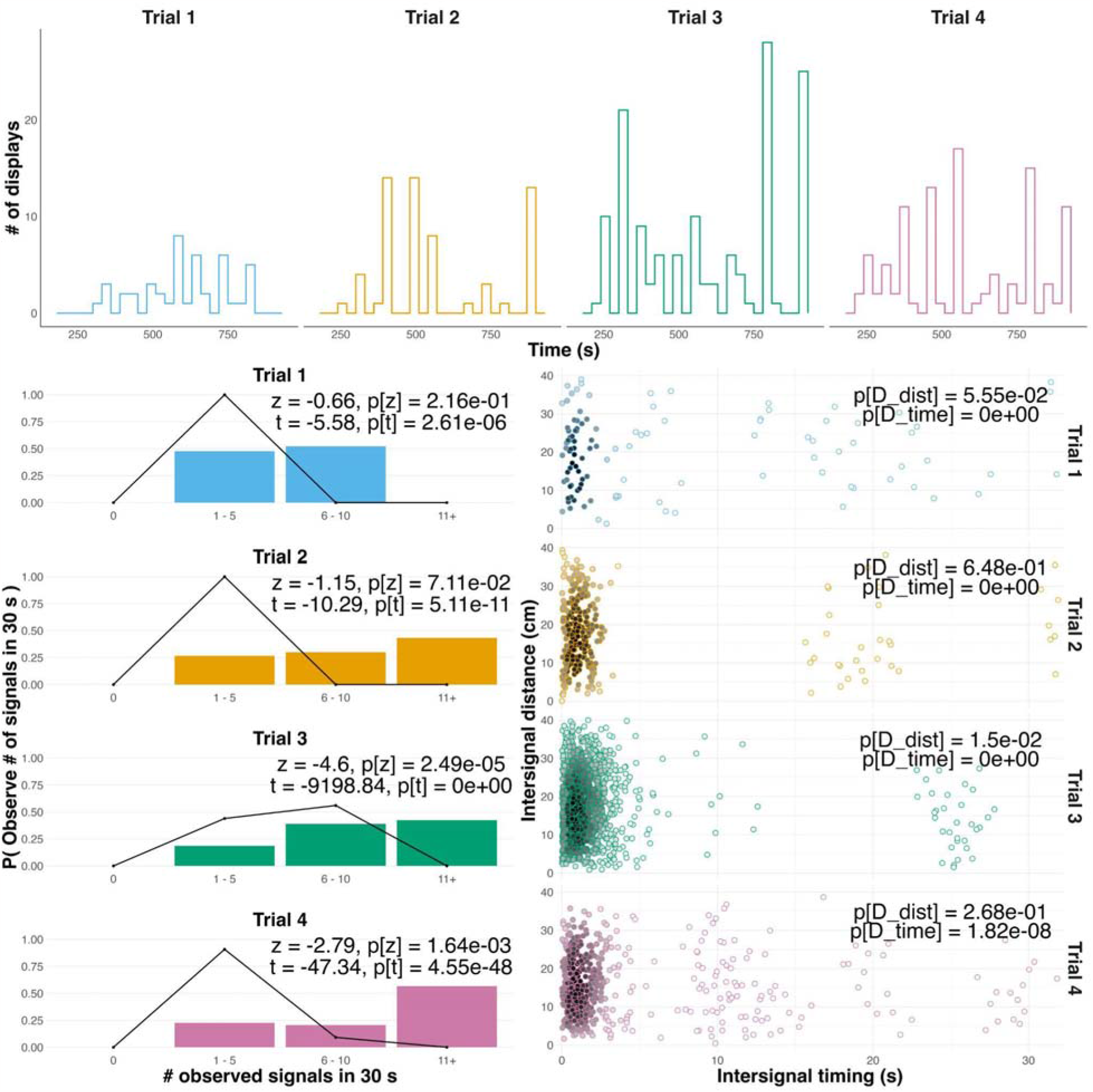
Spatiotemporally annotated sea firefly displays from aquaria during 15 min. observation periods. (A) The number of observed displays (y-axis) over time (x-axis) for 4 separate trials, each with 100 males per tank. (B) Probability of observing N number of signals in 30 s (y-axis; coloured bars) for various signalling rates (x-axis) compared to the null Poisson expectation given the same signalling rate (black line). Goodness-of-fit statistics (Z and T) for a Poisson distribution for each trial, as in [47]. (C) Scatterplot of pairwise signal differences in timing (x-axis) and space (y-axis) show periodic clustering. Comparisons are limited to a maximum of 31 sec, the longest single displays could persist in the environment for a conspecific response (see Table 1). P-values from KS tests comparing the distribution of nearest neighbour display distances (D_dist) and times (D_time) to a null Gaussian distribution. Data coloured by density (darker = more dense).

### Collective behaviour *in situ* is emergent with timescales corresponding to ambient, nightly light cues

Over the course of each night in the grass bed (Fig. 1B), the time between waves increases as minutes past nautical twilight increase (Table S1; data not shown). The signalling niche of ostracods is typically nautical twilight, the time of night after sunset and before moon rise where the brightest natural illumination is starlight [40,45]. Thus, within a night, the number of collective displays peaks ∼45 minutes after sunset (Fig. 1C) and decreases subsequently. The magnitude of this decrease differed between field seasons, likely due to the smaller number of collective events in 2018 compared to 2022 (Table S1).

Between nights, timing between emergent waves is positively correlated with the number of days since the full moon (Fig. 1D, Table S1). As the length of nautical twilight increases during the lunar cycle, the length of time between each subsequent bout of collective signalling also increases. This results in the number of emergent waves decreasing on nights progressively further after the full moon, which again differed between years sampled. Because our sampling window was limited to 1 hour each night, we cannot conclude that the total number of waves during the entire night is greater on days closer to the full moon but we do observe this pattern (Fig. 1D, marginal histogram). Thus we limit our analysis to the rate of collective events in a standardised observation window.

### Individuals produce waves of collective signalling synchronously and nonrandomly

Although apparent to an observer *in situ* (Fig. 1B), we sought to quantify the level of synchrony between displays in a collective. By placing 100 males into aquaria *ex situ* and only tracking their bioluminescent displays, we could observe collective waves emerge in a semi-natural habitat, as seen from the distinct rise and fall in the number of displays over time (Fig. 2A, independent Trails 1 - 4).

Displays co-occur more than would be expected by chance: when compared to a null Poisson distribution of randomly occurring signals at the same mean rate as observed (Fig. 2B, lines), we see that waves have an excess number of signals co-occurring (long-tailed distributions) and a dearth of sampling periods with few signals (Fig. 2B, bars). Using Z and T statistics to compare our observed signal distributions to these Poisson distributions (modified from [47]), we reject the null for all tests except the Z tests for Trials 1 and 2 (Fig. 2B, p-values), which we attribute to the smaller sample sizes of these collectives. Temporal clusters are apparent in the shortest pairwise differences for all displays in a trial (Fig. 2C, but limited to observations within 31 s of each other, the maximum time of any single display Table 1). In all Trials, we reject the null hypothesis that these timing differences come from normal distributions with the same median and variance using Kolmogorov-Smirnov tests (Fig. 2C, x-axis, p[D_time] values). Similar tests for pairwise differences in space detect limited evidence for spatial clustering (Fig. 2C, y-axis, p[D_dist] values) in our tank Trials. Next, we explore potential mechanisms that contribute to this variation in synchrony.

### Collective and individual responses *ex situ* correspond to social stimuli, revealing coordination rules

In *ex situ* experiments, EGD demonstrates differing responses to variation in the duration of incoming stimuli (Fig. 3). Collectively, more animals respond when the stimulus is 4-6 sec long, with fewer responding to stimuli shorter or longer in duration (Fig. 3A, Fig. S4, Table S2). This coincides with the duration of first pulses within individual displays (Table 1). Within two duration experiments, we recorded and measured how individuals time their displays with respect to the beginning and end of the artificial stimuli. Almost no individuals begin displays before stimulus onset (n = 4, data not shown).

**Figure 3.**
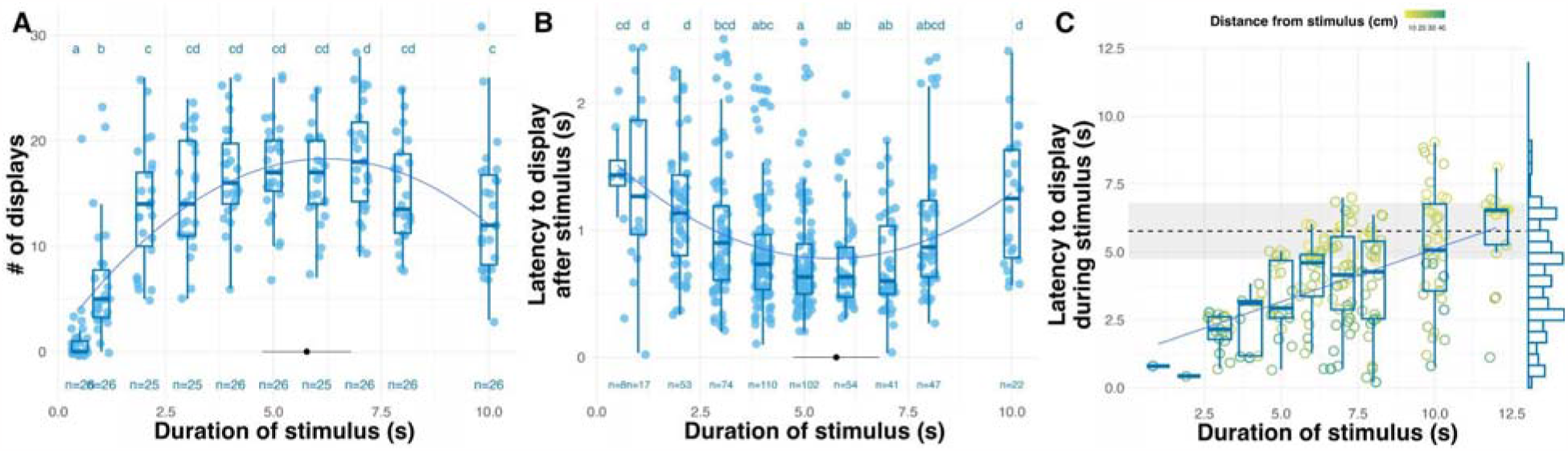
Groups of signalling males vary in the magnitude of their collective (A) and individual (B,C) responses to duration stimuli. (A,B) Boxplots and parabolic curves show that both (A) groups and (B) individuals are most responsive to stimuli lasting 4-6 s. In (A), each datum represents a single experimental trial with multiple males. In (B), each datum is an individual male’s latency to respond after a stimulus presentation. (C) Individuals’ latencies to respond during a stimulus increases with longer stimulus durations (x-axis) and closer to the stimulus (lighter colour = closer). As seen in the marginal plot, males delay their responses only up to ∼7.5 s. In (A,B; datum with whiskers) and (C; dashed line and shaded area), black lines represent the mean + two S.D. for the duration of first pulses from observed displays; see Table 1. For regressions, true model fits for (A-C) in Tables S2, S5, and S4, respectively.

Most males initiate their displays directly after stimulus presentation (n = 700, Chi-squared = 859.79, df = 2, p < 2.2e-16). Limiting our analysis to response latencies of 2.5 s or less (∼ median intersignal time from natural waves, Fig. 2C), we see males respond most quickly after stimuli of intermediate durations (Fig. 3B, Fig. S7, Table S5). Even individuals that begin their displays during a stimulus (n = 197) have increased latencies to initiate a response, both as the stimulus duration increases and the distance from the stimulus decreases (Fig. 3C, Fig. S6, Table S4). However, males only delay up to ∼7.5 s, even if the stimulus duration is greater; again, this coincides well with the maximum duration of first pulses within natural displays.

### Individuals modulate the number of pulses per display based on stimuli distance

Pooling *ex situ* observations (Fig. 4A) and experiments (Fig. 4B, limited to 90 s post-stimulus), we see EGD males farther from a conspecific signal produced displays with more discrete pulses per train. Interdisplay distance describes variation in the timing and latency to display of EGD signals (Fig. 2C, Fig. 3C), as well as the number of pulses within an individual’s display (Fig. 4A, permutation test, p = 0.005; Fig. 4B, permutation test, p = 0), suggesting local interactions are important during collective signalling. The time to the next soonest display (i.e. temporal neighbour) also describes variation in the number of pulses per display (Fig. 4C, Fig. S9A, permutation test, p = 0 ; Fig. S9B, permutation test, p = 0.001). In experimental trials, the distance to the stimulus also influences the number of pulses per display (Fig. S9B, permutation test, p = 0).

**Figure 4.**
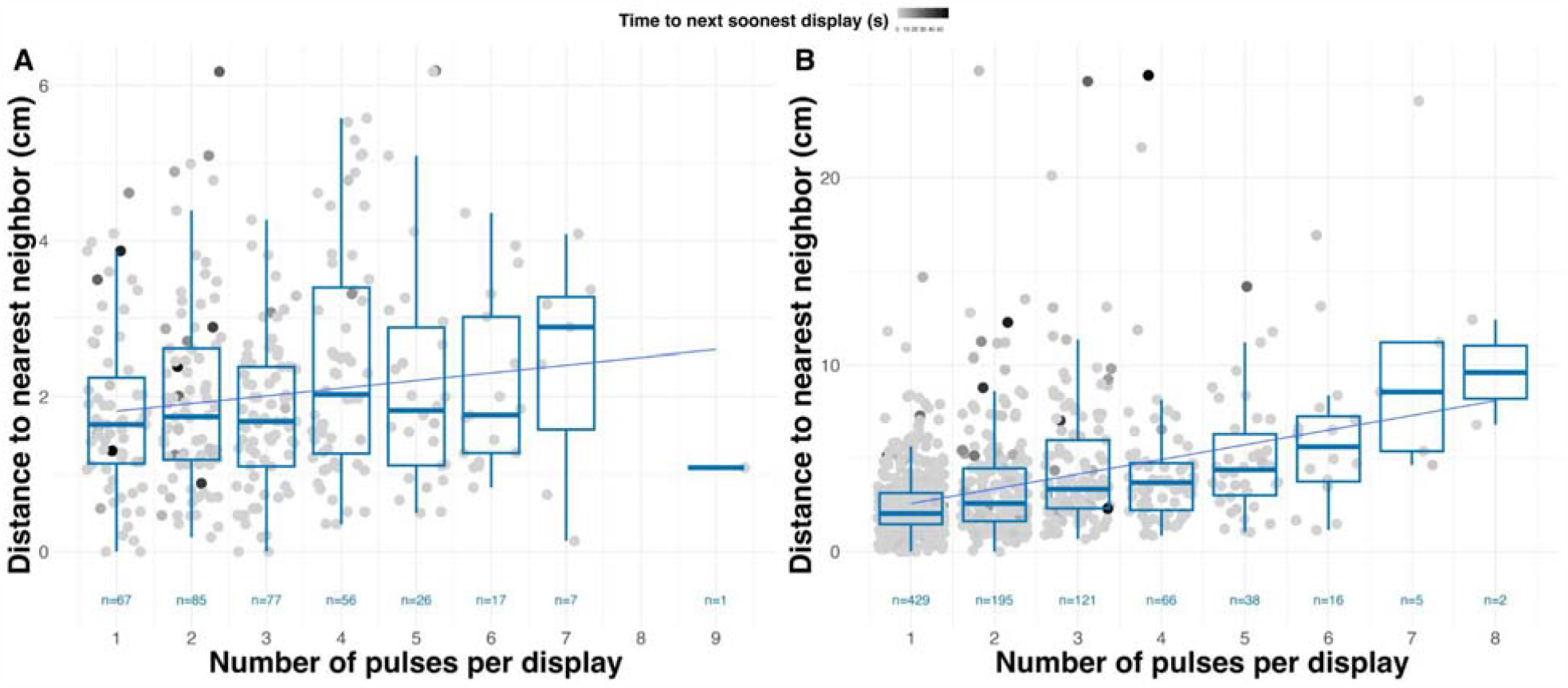
In either (A) observational or (B) experimental trials triggered by an LED stimulus, the distance to the nearest neighbouring male and the time to the next soonest display influence the number of pulses per display. In experimental trials, the distance from the stimulus also influences the number of pulses per display (not indicated). For results of permutation tests, see Fig. S9.

## Discussion

Collective behaviour is sensitive to social conditions and may be constrained by physical or neurological mechanisms. Our data indicate clearly that a recently discovered species of synchronous sea firefly responds to variation in environmental conditions like the duration and intensity of incoming light. Their response magnitude is mediated by variation in these photic stimuli, and other sensory modalities notwithstanding, natural waves of collective behaviour seem to emerge under specific levels of visual stimulation. Our individual-level data reveal that EGD achieves a high level of precision in their signal timing using simple rules: individuals produce bioluminescent signals in response to a visual stimulus, do so most readily integrating light information over time, and have an upper-limit on their response delay. Although variable, sensitivity to external light and intrinsic timing ensures that individuals produce subsequent displays before initiating displays are no longer visible. Understanding variability in these mechanisms will illuminate how synchrony emerges.

Within synchronous waves, there is a lower-limit to the timing between signals, as evidenced by relatively consistent intervals with few to no displays (Fig. 2C). Our field observations (Fig. 1Bii) and *ex situ* data (Fig. 2A, Fig. S2) show wave-like events emerge every ∼30 - 60 s. This pattern is likely influenced by physical constraints on male behaviour, such as swimming invisibly back up in the water column after displaying downwards. However, an intrinsic timing mechanism could also contribute to timing between signalling of different males, like the phase-coupling of oscillators in the rhythmic synchrony of some terrestrial fireflies [12]. Individual latencies to display support the idea that male EGD track temporal changes (Fig. 3B,C), a necessary ingredient for internal phase coupling. However, whether the mechanism underlying the display synchrony we observe in EGD is simply responsive or truely rhythmic requires data on individuals between and during signalling within collectives, and which we could not track in this study due to methodological constraints, but that could be the subject of future studies.

Signalling males seem to plastically adjust their number of pulses in a display as a function of the distance from conspecific displays or light cues, which we predict if bioluminescent trains are signals to conspecifics, and subject to changes in signalling and/or social environment. Plasticity in male mating behaviours is well documented in a congener, whose males change behaviour depending on proximity to a leading display train. Closer individuals switch tactics from luminescent entraining to invisibly sneaking, leading to a reduction in the number of pulses per display due to premature signal termination [44]. In EGD, we expect similar behavioural dynamics are occurring in these spree leks. Individuals farther from one another or from a stimulus create displays with more pulses, suggesting that closer males could be rapidly switching tactics from entraining to sneaking. Conversely, individuals may be aborting their signals earlier based on local timing cues from neighbours who produce displays with few pulses.

We do not find behavioural sensitivity to neighbour proximity in the spacing of individual displays within *ex situ* collectives (Fig. 2C). This may be the result of a relatively unobstructed visual space, allowing individuals to simply synchronise with visual neighbours of varying distance, but with finer scale interactions modulated by changes in intensity. Visual obstructions may introduce spatial constraints to collective behaviour, as in terrestrial fireflies [27], by reducing incident light intensity.

Collective behaviour is also sensitive to ecological cues. The decrease in the frequency and number of collective signalling events in EGD across a lunar cycle supports the hypothesis that the available amount of time of complete darkness during the night is a valuable ecological resource [45]. Other luxorine ostracods in sympatry occupy distinct temporal niches throughout the night [40], suggesting some timing mechanism helps each regulate their peak display window. Within *P. annecohenae*, there were no patterns of display activity related to long-term lunar cycles; signalling behaviour in the wild was best predicted by overall light level, which co-varies with the lunar cycle [45]. In ostracods, darkness may coincide with other cyclical processes like availability of newly receptive females. Given our results, EGD seems sensitive to short-timescale light levels, similar to *P. annecohenae*, because we were able to manipulate their propensity to display *ex situ* by keeping tanks illuminated despite the time of night or lunar phase. Fireflies behave similarly during reductions in daylight (e.g. solar eclipse [48]). However, there may be a circalunar cycle that is important to the timing of EGD reproductive behaviours, which is common in other marine organisms (e.g. corals, spawning annelids, etc.), and which we are not able to rule out [49]. Manipulative experiments could demonstrate the effects of light sensitivity and intrinsic, clock-like behavioural mechanisms on behaviour, and if they contribute to behavioural or community diversity.

For luminescent organisms ambient light is also a valuable resource - in this case, a signal of their social niche. Correctly identifying and evaluating the social salience of available light levels in the environment poses a challenge. Regardless of modality, filtering signals from noise is a sensory and perceptual task [50] that has large fitness consequences (e.g. [51,52]). EGD may have adaptations to differentiate low-levels of ambient light from dim but increasingly bright conspecific displays. Terrestrial firefly species adaptively vary their peak spectral emission (≈ colour) to increase visibility via contrast enhancement as background light changes during night [53,54]. But unlike insects, ostracods vary little in their peak spectral emission [39], possibly due to constraints of signalling in the marine environment, which selects for a narrower range of wavelengths in the visible spectrum [55] than terrestrial habitats. To increase signal recognition, one adaptive strategy may be in varying mechanisms of temporal integration, producing bandpass filters that could ignore changes in light intensity outside a particular rate. Our data show that EGD are attuned to temporal features of stimuli (Fig. 3). Costs of recognition errors may be higher in the temporal domain given the prevalence of other non-salient light cues in the environment such as moonlight, the bioluminescent signals of other organisms (dinoflagellates, cnidarian polyps, syllid worms [56]), and occasionally other ostracod species that occur where two habitat margins overlap (pers. obs), offering a larger opportunity for reproductive interference.

Collective signals are known in arthropods, but their adaptive functions can be hard to assess experimentally. Some success has been made in fiddler crabs [57] and bees [58], using mathematical models of synchronous insects [59], and recently in a non-arthropod, the sulphur molly [60]. In synchronous fireflies, males should benefit from collective signals because females respond more [61]; therefore synchrony may aid in species recognition by decreasing visual clutter. However, in fiddler crabs [57] and katydids [59], females prefer leading males, producing increased competition between faster males - thus sexual selection via female precedence preference results in incidental synchrony (an “epiphenomenon”). Following males still accrue potential mates [62]. These results align with models of competitive interactions between signallers that produce collective dynamics [19,59]. In the case of luminous ostracods like EGD, sensitivity to neighbouring signals may be the target of sexual selection. Synchrony could also be an example of bet hedging costs to individual recognition during mating in the face of predation. Although we cannot rule out this alternative currently, bioluminescence in ostracods is believed to be anti-predatory [31]. Which of these three hypotheses is most explanatory requires manipulative experiments to assay the costs and benefits of different behaviours. Marine ostracods like EGD have a number of unique attributes that make them tractable for future work: (1) they are abundant, small, and non-seasonal; (2) sexes, lifestages, and signals are easy to differentiate; and (3) both groups and individuals will behave in captivity.

Here, we describe a unique marine system that is seemingly convergent with terrestrial fireflies in the emergence of synchronous mating signals. Like their aerial arthropod cousins, sea fireflies produce individual luminous mating displays that coalesce into collectives based on local rules of interaction. One assumption in investigating synchronous mating signals using a comparative framework is that these emergent collective phenomena are heritable. It is difficult to hypothesise about the genetic basis of synchrony (although some work attempts to find neuronal or genetic mechanisms, as referenced in [63]). Rather than solely emergent from plastic responses to the social or ecological environment, variation in the behavioural rules that underlie the emergence of synchrony in each taxon should be transmissible between generations, like sensitivity to conspecific signals, and are both quantifiable and can be linked to the collective properties. These types of behaviours seem more tractable for understanding how synchronous mating systems repeatedly evolve. Being clear about the behavioural mechanisms that produce synchrony in diverse clades, and measuring analogous variables across clades in similar ways will be key in making studies comparable in a phylogenetic framework to understand the evolutionary pressures that produce emergent behaviour [5,64,65].

## Methods

### A comment on synchrony versus entrainment in Ostracoda

Although previous work documents spatial clustering with strong temporal entrainment in some ostracod species [43,66], synchrony has never been described. Synchrony is a pattern distinct from an entrainment process, but the two are often conflated [67]. In chronobiology, entrainment is when an internal, rhythm-generating oscillator becomes phase coupled to another cyclical process (i.e. tidal, circadian, another organism’s oscillatory mechanism, etc.), and is sometimes called “rhythmic synchrony”. Previous descriptions of ostracod signalling describe entrainment as “loose synchrony” [37,44], seemingly invoked to provide a logical comparison with other synchronous displays like in terrestrial fireflies [68]. However, distinguishing between synchronous behaviours due to an entrained, clock-like mechanism or non-entraining responsiveness (i.e. “simple synchrony”) requires manipulative studies that have not been performed within luxorine ostracods. Previous attempts to elicit single displays from individuals in lab settings have failed as males seem to require the presence of congeners [44]. From these studies in a non-synchronous species, we see interdisplay intervals ∼19 s in groups of four signalling males, but we lack evidence of individual rhythmicity [38]. In this study, we attempt to strike a balance between rhythmic and simple synchrony, but primarily deal with simple synchrony despite the apparent relevance and previous invocations of entrainment in this system.

### Animal collection and maintenance

We collected both male and female *Photeros* sp. “EGD” (nightly from 11 - 23 May 2017 and 21 - 26 September 2018, respectively) using baited conical traps [69] placed in the grass beds of Punto Manglar (9.332464, -82.254105) off the coast of Bocas del Toro, Panama. Males were also collected via sweep netting through visible displays. Adult animals were sorted by eye the next day and kept in tupperware with fresh sea water (Gladware 24oz. containers) in small batches. Each day we checked the health of individuals and changed their sea water; we fed them fish flakes (Seachem NutriDiet MarinePlus Enhanced Marine Flakes with Probiotics) every other day.

### Characterising individual EGD displays

EGD displays were measured using two separate systems. Timing characteristics were collected from continuous filming of collective displays during field work in May 2017. Underwater scenes were filmed with a stabilised Sony A7 and Atomos Shogun system in custom underwater housing, as in [70]. From single video, we then measured the duration, interpulse interval, total number of pulses, and brightness (see below) from displays that were contemporaneous but distinct.

Distance measures were collected from continuous filming of collective displays during field work in September 2018. Using two WATEC 910HX cameras at a fixed distance from one another in a custom underwater housing and that simultaneously record the same vantage, we could calculate the absolute distances (Oakley et al. *in prep*.) [71]. From this video, we measured the interpulse distances and total display length.

To measure the intensity of pulses, we used two methods. First, we placed multiple individuals in a glass aquarium and recorded their displays using a Hamatsu photomultiplier tube (PMT) placed perpendicular to and at a fixed distance from the tank. From intensity over time recordings, we isolated trains of peaks that were discrete, decayed continuously without interruption, were grouped in time, and non-overlapping, matching expectations from similar recordings of another species [38]. In a separate analysis, we took fixed video from underwater scenes (as above) and used ImageJ to isolate the pixels of both single pulses within trains and a similar area of video without pulses outside of trains. We subtracted this background pixel intensity from the peak pulse pixel intensity for every frame. For data collected with either method, we normalised all pulse values within a train to the first pulse. Although pulse intensities are relative within a recording, reducing variation due to methodology, overall differences in sensitivity and distance from the detector may influence the ability to discriminate pulses from background noise.

### *In situ* observations

Field observations occurred every 2 - 3 nights after the full moon during 27 September - 10 October 2018 and 13 - 24 July 2022. Directly after the onset of nautical twilight and the first visible wave of collective bioluminescence, a single observer stood in the seagrass and counted the number of events during 60 min in a 2 m radius from the observer’s field of vision. Displays were considered collective waves if 5 or more individual displays were visible within ∼5 s of one another. Within a field season, the observer made recordings from the same place in the seagrass bed and facing the same cardinal direction.

### *Ex situ* observation and experiments

Experiments occurred nightly from 18 - 22 May 2017 between 2100 hrs and 0300 hrs in water tables. During the day, 100 male EGD were checked by eye and separated into batches for experiments that night. At ∼1800 hrs, we filled tanks (100 x 100 cm) with clean sea water to a depth of 15-20 cm, placing each batch of males into a separate tank, and using up to 4 tanks per night. To prevent males from signalling and to simulate the onset of night, we turned on two puck lights set on the edge of each tank to shine white LEDs into the tanks. When we were ready, lights would be extinguished, animals were given 20 min. to adapt to darkness, and we observed tanks for 15 minutes before exposing them to any stimulus. Each day after experiments from the night before, tanks were drained and washed with fresh water to remove any animals, and then allowed to air dry.

For stimulus exposure experiments (both intensity and duration), we used a dive light with white LED lights covered with a blue film as the stimulus, which was the only light source available.

Anecdotally, this light source was used to trigger waves of male displays *in situ*, and thus was deemed adequate to elicit a response from males in our tanks. We connected the LED to an Arduino mini-controller and pre-programmed a range of square-pulse signals to drive the LED on or off. We used a nested design to expose each tank every night tested (deemed a trial) to the full range of conditions within a stimulus type three times each. During each trial, we randomly assigned the order of the test conditions within a tank, and between trials, we randomly assigned the order of testing among tanks. We used Excel to generate four series of random sequences of stimulus conditions and randomly assigned one series to each tank every night. We programmed these four series into the Arduino code to loop automatically through each stimulus within a series at the initiation of each trial. An observer recorded the number of animals that responded to a stimulus by counting the number of unique displays in a tank within 1 minute of the stimulus onset. Stimuli durations were 500 ms, and 1, 2, 3, 4, 5, 6, 7, 8, 10 s long.

For two tank experiments and four 15-min observation trials (Trials 1 - 4), we also filmed the tanks obliquely from above, approximately overhead, using a Sony A7S with High ISO setting and attached to an Atomos Shogun for video recording in 4K. Videos were exported to Final Cut Pro, and then observers used the software Tracker [72] to record the start time, position of first pulse, and number of pulses per display during the trials. In the two experimental trials, observers also noted the relative timing of display start times with respect to stimulus start and end time, and total duration.

### Data analysis

Almost all statistical analyses were performed using linear models unless otherwise noted. Specifics for each model are below and can be found in the code (Electronic Supplementary Material). For all our models where possible, we used the ‘check_model’ function from the package “performance” to visually inspect residuals and check model assumptions. If we tested the presence of interactions or other variables that were subsequently excluded because they were not significant, we used the ‘anova’ function to compute a chi-squared test and compare the simple model versus the more complex model to decide if explanatory variables should be retained. All data and statistical analyses were performed in R (vers. 3.6.2) with RStudio (vers. 2022.12.0+353). Figure colours from [73,74]. In all scatterplots, data are jittered for visualisation purposes, and all regression lines are only demonstrative unless specifically noted. Model fits can be found in the supplementary data.

To understand how different timescales influence the observed number of collective displays, we fit a linear model using OLS of the following construction:

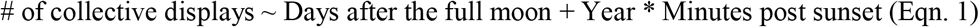

Results in Table S1.

To test if the number of responding displays differed among experimental treatments, we used the following models fit with OLS. Although our data are counts, they reasonably met the assumptions of a normal distribution to use linear models:

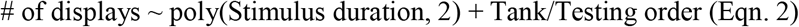

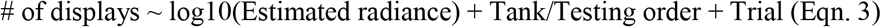

We nested the order in which tanks were tested within tank identity because we could not test every tank across every level of the testing order. We also included a trial variable to capture variation due to the stimulus presentation order, which was randomised within trials but the same between trials of the same type (see Methods above). Results in Table S2 and S3, respectively.

To test if individual latency to display differed among experimental treatments, we used the following models fit with OLS:

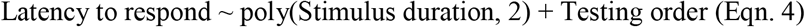

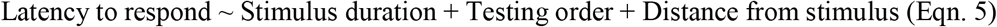

Results in Tables S4 and S4, respectively. For results from Eqn. 2 - 4, we used the estimated marginal means to look for pairwise group differences, as denoted by the letters for each group. Groups that are not statistically different after a Sidak method correction for multiple comparisons share the same label.

To show that signals are clustered in time (i.e. synchronised) and/or space, we used four approaches. First, we measured pairwise differences between signals for their timing and their spatial distance during each trial. We limited our analysis to signals produced at most within 31 s of one another because this is the maximum duration of individual signals produced in the wild (Table 1), and therefore relevant to the response time of other individuals. We expect to see discrete as opposed to continuous distributions of pairwise differences if signals are synchronised and/or coincident. Conversely, if signals are randomly generated over time or space, pairwise differences between signal occurrences should approximate a normal distribution (limited to 0 minimum). To test this pattern, for each trial we generated a null dataset using a truncated Normal distribution with the same median and standard deviation as the observed data, and used Kolmogorov-Smirnov tests to compute p-values.

Next, to statistically test if displays are synchronised, we compared the expected number of displays per sampling time to the observed number of displays, as from [47]. Because displays are discrete events, we can consider them to occur from a Poisson process. For a given trial, we used the total number of observed displays divided by the total observation time to calculate the given display rate. We used this rate to simulate a null expectation asking how much of our sampling time (in 30 s bins) contains 0,1,2 … *n* or more displays. We then compared the expected probability of seeing *n* or more displays in 30 s (Fig. 4B, black line) with our observations (Fig. 4B, coloured bars).

Lastly, we used two statistical tests from [47]. Briefly, these statistics are goodness-of-fit tests to compare an observed distribution to that of a Poisson distribution that assumes recorded occurrences (i.e. displays) are uncorrelated. Because our data are coded at the level of single displays (each composed of many pulses), and not at the level of flashes within bursts (the analogous situation in terrestrial fireflies, and for which data these tests were devised), we modified the T test to exclude excess correlation in the structure of measured occurrences (see code in ESM). Significant p-values calculated from these tests would reject the null hypothesis that the observed distribution was generated from a Poisson process.

To understand the factors affecting variation in the number of pulses per display, the residuals in our data did not meet any normality distribution assumptions in order to use linear models. As such, we used a permutation approach to understand the effect of three variables of interest. We performed 1000 permutations per predictor variable, permuting the pairwise distance matrix for each predictor within trials (observational, Fig. 6A, Fig. S9A using Eqn 6.; or experimental; Fig. 6B, Fig. S9B using Eqn 7.) and using the following models to regress the variables against pulse number:

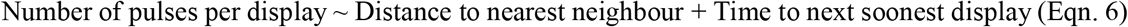

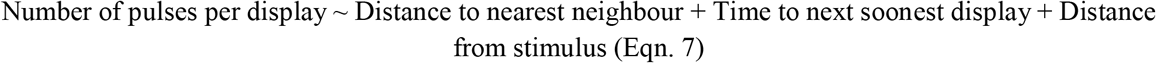

Because this is computationally intensive, we limited our analysis to understanding these three predictor variables. A caveat to the interpretation of these results is that all these metrics have some level of correlation with one-another (Fig. S10, S11), although our permutation test results are robust to such correlations by design. To generate p-values, we compared the number of t-statistics from the distribution of t-statistics from the permuted linear regressions that were more extreme than our observed statistic, which should have a consistent Type I error rate [75].

The following R packages were used to handle, analyse, and visualise all data: tidyverse [76], ggplot2 [77], readxl [78], RColorBrewer [79], ggpmisc [80], ggpubr [81], dplyr [82], ggExtra [83], MASS [84], ggpointdensity [85], tidyr [86], patchwork [87], png [88], spatstat [89], broom [90], stringr [91], ggbreak [92], performance [93], sjPlot [94], jtools [95], emmeans [96], multcomp [97], multcompView [98], scales [99], GGally [100]. See electronics supplementary material for details.

## Supporting information

Supplemental Results, Figures, and Tables

## Acknowledgements

Thanks to Emily A. Ellis, Vanessa Gonzalez, Nick Reda, Wesley Peery, Cheyenne McKinley, and Rebecca M. Varney for field support. And to Daniel Escobar Camacho for measuring the intensity and emission spectra of our *ad hoc* field stimulus. We thank Drs. Rachel Collins and Harilaos Lessios at the Smithsonian Tropical Research Institute (STRI) for their support, as well as members of the STRI Bocas del Toro Field Station, especially Plinio Gondola, Marcos Alvarez, and Felicito Caito for their assistance in completion of this project. We appreciate advice from Elizabeth Hobson on various statistical analysis.

## Funding statement

NMH was supported by the NSF GRFP, NSF PRFB, and a STRI Short-term Fellowship; he received a Sigma Xi Grants in Aid of Research (G20141015722209), a block grant from the Department of Ecology, Evolution, and Marine Biology at UCSB, and a Lerner-Grey Grant for Marine Research from the American Museum of Natural History to partially fund this work. GAG was supported by NSF (#DEB-1457754); THO was supported by NSF (#DEB-1457439). All work was conducted under the auspices of permits obtained from the Panamanian government and provided by MiAmbiente (Permits #SE/A-54-18, #SEX/A-78-18, #ARB-082-2022).

## Notes

https://github.com/NikoHensley/ESM_egd_collective_behavior

### Competing Interest Statement

The authors have declared no competing interest.

### Summary of Updates

In accordance with peer review, we have updated some of the clarity of text, and added more statistical tests. We have also more appropriately attributed effort to a new co-author.

## References

1. Shishkov O, Peleg O. 2022 Social insects and beyond: The physics of soft, dense invertebrate aggregations. Collective Intelligence 1, 26339137221123758.

2. Sumpter DJT. 2010 Collective Animal Behavior. Princeton University Press.

3. Ballerini M et al. 2008 Interaction ruling animal collective behavior depends on topological rather than metric distance: evidence from a field study. Proc. Natl. Acad. Sci. U. S. A. 105, 1232–1237.

4. Evangelista DJ, Ray DD, Raja SK, Hedrick TL. 2017 Three-dimensional trajectories and network analyses of group behaviour within chimney swift flocks during approaches to the roost. Proc. Biol. Sci. 284. (doi:10.1098/rspb.2016.2602)

5. Gordon DM. 2014 The ecology of collective behavior. PLoS Biol. 12, e1001805.

6. Gordon DM. 2021 Measuring collective behavior: an ecological approach. Theory Biosci. 140, 353–360.

7. King AJ, Fehlmann G, Biro D, Ward AJ, Fürtbauer I. 2018 Re-wilding Collective Behaviour: An Ecological Perspective. Trends Ecol. Evol. 33, 347–357.

8. Sheppard LW, Mechtley B, Walter JA, Reuman DC. 2020 Self-organizing cicada choruses respond to the local sound and light environment. Ecol. Evol. 10, 4471–4482.

9. Ouellette NT, Gordon DM. 2021 Goals and Limitations of Modeling Collective Behavior in Biological Systems. Frontiers in Physics 9. (doi:10.3389/fphy.2021.687823)

10. Dibnah AJ, Herbert-Read JE, Boogert NJ, McIvor GE, Jolles JW, Thornton A. 2022 Vocally mediated consensus decisions govern mass departures from jackdaw roosts. Curr. Biol. 32, R455– R456.

11. Garnier S, Gautrais J, Theraulaz G. 2007 The biological principles of swarm intelligence. Swarm Intelligence 1, 3–31.

12. Acebrón JA, Bonilla LL, Pérez Vicente CJ, Ritort F, Spigler R. 2005 The Kuramoto model: A simple paradigm for synchronization phenomena. Reviews of Modern Physics. 77, 137–185. (doi:10.1103/revmodphys.77.137)

13. Araujo SBL, Rorato AC, Perez DM, Pie MR. 2013 A spatially explicit model of synchronization in fiddler crab waving displays. PLoS One 8, e57362.

14. Rathore A, Isvaran K, Guttal V. 2023 Lekking as collective behaviour. Philos. Trans. R. Soc. Lond. B Biol. Sci. 378, 20220066.

15. Buck J, Buck E. 1978 Toward a Functional Interpretation of Synchronous Flashing by Fireflies. Am. Nat. 112, 471–492.

16. Legett HD, Page RA, Bernal XE. 2019 Synchronized mating signals in a communication network: the challenge of avoiding predators while attracting mates. Proc. Biol. Sci. 286, 20191067.

17. Greenfield MD, Aihara I, Amichay G, Anichini M, Nityananda V. 2021 Rhythm interaction in animal groups: selective attention in communication networks. Philos. Trans. R. Soc. Lond. B Biol. Sci. 376, 20200338.

18. Greenfield MD. 2005 Displays in Arthropods and Anurans. Adv. Stud. Behav. 35, 1.

19. Greenfield MD, Tourtellot MK, Snedden WA. 1997 Precedence effects and the evolution of chorusing. Proceedings of the Royal Society of London. Series B: Biological Sciences 264, 1355–1361.

20. McLellan CF, Montgomery SH. 2023 Towards an integrative approach to understanding collective behaviour in caterpillars. Philos. Trans. R. Soc. Lond. B Biol. Sci. 378, 20220072.

21. Ramaswamy K, Cocroft RB. 2009 Collective signals in treehopper broods provide predator localization cues to the defending mother. Anim. Behav. 78, 697–704.

22. Hartbauer M, Siegert ME, Fertschai I, Römer H. 2012 Acoustic signal perception in a noisy habitat: lessons from synchronising insects. J. Comp. Physiol. A Neuroethol. Sens. Neural Behav. Physiol. 198, 397–409.

23. Nityananda V, Balakrishnan R. 2021 Synchrony of complex signals in an acoustically communicating katydid. J. Exp. Biol. (doi:10.1242/jeb.241877)

24. Backwell P, Jennions M, Passmore N, Christy J. 1998 Synchronized courtship in fiddler crabs. Nature 391, 31–32.

25. Backwell PRY. 2019 Synchronous waving in fiddler crabs: a review. Curr. Zool. 65, 83–88.

26. Buck J, Buck E. 1976 Synchronous fireflies. Sci. Am. 234, 74–9, 82–5.

27. Sarfati R, Hayes JC, Peleg O. 2021 Self-organization in natural swarms of Photinus carolinus synchronous fireflies. Sci Adv 7. (doi:10.1126/sciadv.abg9259)

28. Buck J. 1988 Synchronous rhythmic flashing of fireflies. II. Q. Rev. Biol. 63, 265–289.

29. Buck J, Buck E. 1968 Mechanism of rhythmic synchronous flashing of fireflies. Fireflies of Southeast Asia may use anticipatory time-measuring in synchronizing their flashing. Science 159, 1319–1327.

30. Morin JG. 1986 Firefleas of the Sea: Luminescent Signaling in Marine Ostracode Crustaceans. Fla. Entomol. 69, 105–121.

31. Ellis EA et al. 2022 Sexual signals persist over deep time: ancient co-option of bioluminescence for courtship displays in cypridinid ostracods. Syst. Biol. (doi:10.1093/sysbio/syac057)

32. Brock TD. 1970 High Temperature Systems. Annu. Rev. Ecol. Syst. 1, 191–220.

33. Vandekerkhove J, Martens K, Rossetti G, Mesquita-Joanes F, Namiotko T. 2013 Extreme tolerance to environmental stress of sexual and parthenogenetic resting eggs ofEucypris virens(Crustacea, Ostracoda). Freshw. Biol. 58, 237–247.

34. Oakley TH. 2005 Myodocopa (Crustacea: Ostracoda) as models for evolutionary studies of light and vision: multiple origins of bioluminescence and extreme sexual dimorphism. Hydrobiologia 538, 179–192.

35. Cohen AC, Morin JG. 2003 Sexual Morphology, Reproduction and the Evolution of Bioluminescence in Ostracoda. The Paleontological Society Papers 9, 37–70.

36. Morin JG. 2019 Luminaries of the reef: The history of luminescent ostracods and their courtship displays in the Caribbean. J. Crustacean Biol. 39, 227–243.

37. Morin JG, Cohen AC. 2010 It’s All About Sex: Bioluminescent Courtship Displays, Morphological Variation and Sexual Selection in Two New Genera of Caribbean Ostracodes. J. Crustacean Biol. 30, 56–67.

38. Rivers TJ, Morin JG. 2008 Complex sexual courtship displays by luminescent male marine ostracods. J. Exp. Biol. 211, 2252–2262.

39. Hensley NM et al. 2021 Selection, drift, and constraint in cypridinid luciferases and the diversification of bioluminescent signals in sea fireflies. Mol. Ecol. 30, 1864–1879.

40. Gerrish GA, Morin JG. 2016 Living in sympatry via differentiation in time, space and display characters of courtship behaviors of bioluminescent marine ostracods. Mar. Biol. 163, 190.

41. Rivers TJ, Morin JG. 2013 Female ostracods respond to and intercept artificial conspecific male luminescent courtship displays. Behav. Ecol. 24, 877–887.

42. Torres E, Morin JG. 2007 Vargula Annecohenae, a New Species of Bioluminescent Ostracode (Myodocopida: Cypridinidae) from Belize. J. Crustacean Biol.

43. Cohen AC, Morin JG. 2010 Two New Bioluminescent Ostracode Genera, Enewton And Photeros (Myodocopida: Cypridinidae), with Three New Species from Jamaica. J. Crustacean Biol. 30, 1–55.

44. Rivers TJ, Morin JG. 2009/9 Plasticity of male mating behaviour in a marine bioluminescent ostracod in both time and space. Anim. Behav. 78, 723–734.

45. Gerrish GA, Morin JG, Rivers TJ, Patrawala Z. 2009 Darkness as an ecological resource: the role of light in partitioning the nocturnal niche. Oecologia 160, 525–536.

46. Torres E, Cohen AC. 2005 Vargula morini, a new species of bioluminescent ostracode (Myodocopida: Cypridinidae) from Belize and an associated copepod (Copepoda: Siphonostomatoida: Nicothoidae). J. Crustacean Biol. 25, 11–24.

47. Sarfati R, Gaudette L, Cicero JM, Peleg O. 2022 Statistical analysis reveals the onset of synchrony in sparse swarms of Photinus knulli fireflies. J. R. Soc. Interface 19, 20220007.

48. Branham MA, Faust LF. 2019 Firefly Flashing Activity during the Totality Phase of a Solar Eclipse. entn 128, 191–203.

49. Raible F, Takekata H, Tessmar-Raible K. 2017 An Overview of Monthly Rhythms and Clocks. Front. Neurol. 8, 189.

50. Pollack L, Munson A, Savoca MS, Trimmer PC, Ehlman SM, Gil MA, Sih A. 2022 Enhancing the ecological realism of evolutionary mismatch theory. Trends Ecol. Evol. 37, 233–245.

51. Senzaki M et al. 2020 Sensory pollutants alter bird phenology and fitness across a continent. Nature 587, 605–609.

52. Maan ME, Seehausen O, Groothuis TGG. 2017 Differential Survival between Visual Environments Supports a Role of Divergent Sensory Drive in Cichlid Fish Speciation. Am. Nat. 189, 78–85.

53. Hall DW, Sander SE, Pallansch JC, Stanger-Hall KF. 2016 The evolution of adult light emission color in North American fireflies. Evolution 70, 2033–2048.

54. Lall AB, Seliger HH, Biggley WH, Lloyd JE. 1980 Ecology of colors of firefly bioluminescence. Science 210, 560–562.

55. Widder EA. 2010 Bioluminescence in the ocean: origins of biological, chemical, and ecological diversity. Science 328, 704–708.

56. Morin JG. 1983 Coastal Bioluminescence: Patterns and Functions. Bull. Mar. Sci. 33, 787–817.

57. Kahn AT, Holman L, Backwell PRY. 2014 Female preferences for timing in a fiddler crab with synchronous courtship waving displays. Anim. Behav. 98, 35–39.

58. Nguyen DMT, Iuzzolino ML, Mankel A, Bozek K, Stephens GJ, Peleg O. 2021 Flow-mediated olfactory communication in honeybee swarms. Proc. Natl. Acad. Sci. U. S. A. 118. (doi:10.1073/pnas.2011916118)

59. Greenfield MD, Roizen I. 1993 Katydid synchronous chorusing is an evolutionarily stable outcome of female choice. Nature 364, 618–620.

60. Poel W, Daniels BC, Sosna MMG, Twomey CR, Leblanc SP, Couzin ID, Romanczuk P. 2022 Subcritical escape waves in schooling fish. Sci Adv 8, eabm6385.

61. Moiseff A, Copeland J. 2010 Firefly synchrony: a behavioral strategy to minimize visual clutter. Science 329, 181.

62. Harrison LM, Melo GC, Perez DM, Backwell PRY. 2021 Why signal if you are not attractive? Courtship synchrony in a fiddler crab. Behav. Ecol. 32, 1224–1229.

63. Greenfield MD, Honing H, Kotz SA, Ravignani A. 2021 Synchrony and rhythm interaction: from the brain to behavioural ecology. Philos. Trans. R. Soc. Lond. B Biol. Sci. 376, 20200324.

64. Doyle JC, Csete M. 2011 Architecture, constraints, and behavior. Proc. Natl. Acad. Sci. U. S. A. 108 Suppl 3, 15624–15630.

65. Papadopoulou M, Fürtbauer I, O’Bryan LR, Garnier S, Georgopoulou DG, Bracken AM, Christensen C, King AJ. 2023 Dynamics of collective motion across time and species. Philos. Trans. R. Soc. Lond. B Biol. Sci. 378, 20220068.

66. Cohen AC, Morin JG. 1986 Three new luminescent ostracodes of the genus Vargula (Myodocopida, Cypridinidae) from the San Blas region of Panama. Natural History Museum of Los Angeles County.

67. Bittman EL. 2021 Entrainment Is NOT Synchronization: An Important Distinction and Its Implications. J. Biol. Rhythms 36, 196–199.

68. Carlson AD, Copeland J. 1985 Flash Communication in Fireflies. Q. Rev. Biol. 60, 415–436.

69. Cohen AC, Oakley TH. 2017 Collecting and processing marine ostracods. J. Crustacean Biol. 37, 347–352.

70. Hensley Nicholai M., Ellis Emily A., Gerrish Gretchen A., Torres Elizabeth, Frawley John P., Oakley Todd H., Rivers Trevor J. 2019 Phenotypic evolution shaped by current enzyme function in the bioluminescent courtship signals of sea fireflies. Proceedings of the Royal Society B: Biological Sciences 286, 20182621.

71. Oakley TH, Motta CA, Saha R, Locker-Cameron TR, Hensley NH, Rivers TJ. 2018 Waterborne Autonomous Low Light Electrostereovideography (WALL-E) to Quantify Luminous Courtship Signals of Ostracods. In INTEGRATIVE AND COMPARATIVE BIOLOGY, pp. E389–E389. OXFORD UNIV PRESS INC JOURNALS DEPT, 2001 EVANS RD, CARY, NC 27513 USA.

72. Brown D, Cox AJ. 2009 Innovative Uses of Video Analysis. Phys. Teach. 47, 145–150.

73. Wilke CO. 2019 Fundamentals of Data Visualization: A Primer on Making Informative and Compelling Figures. ‘O’Reilly Media, Inc.’

74. Okabe M, Ito K. In press. Color universal design (cud)-how to make figures and presentations that are friendly to colorblind people. Retrieved April

75. Anderson MJ, Robinson J. 2001 Permutation tests for linear models. Aust. N. Z. J. Stat. 43, 75–88.

76. Wickham H, Wickham MH. In press. Package tidyverse. the ‘Tidyverse

77. Wickham H. 2011 Ggplot2. Wiley Interdiscip. Rev. Comput. Stat. 3, 180–185.

78. Wickham H, Bryan J. In press. readxl: Read excel files. R package version

79. Neuwirth E, Neuwirth ME. 2014 Package ‘RColorBrewer’. ColorBrewer Palettes

80. Aphalo PJ. In press. ggpmisc: Miscellaneous Extensions to ‘ggplot2’. R package version 0.3

81. Kassambara A. In press. ggpubr:’ggplot2’ based publication ready plots (Version 0.1. 7). desde https://CRAN.R-project.org/package=ggpubr

82. Wickham H, François R, Henry L, Müller K. In press. dplyr: A grammar of data manipulation. R package version 0.4

83. Attali D, Baker C. In press. ggExtra: Add marginal histograms to ‘ggplot2’, and more ‘ggplot2’enhancements. R package version

84. Ripley B, Venables B, Bates DM, Hornik K, Gebhardt A, Firth D, Ripley MB. 2013 Package ‘mass’. Cran r 538, 113–120.

85. Kremer LPM. 2019 ggpointdensity: A cross between a 2D density plot and a scatter plot.

86. Wickham H, Wickham MH. 2017 Package ‘tidyr’. Easily Tidy Data with’spread’and’gather ()’Functions

87. Pedersen TL. 2019 Package ‘patchwork’. R package http://CRAN.R-project.org/package=patchwork.Cran

88. Urbanek S. In press. png: Read and write PNG images. R package version 0. 1-7

89. Baddeley A, Turner R. 2005 spatstat: An R Package for Analyzing Spatial Point Patterns. J. Stat. Softw. 12, 1–42.

90. Robinson D. 2014 broom: An R Package for Converting Statistical Analysis Objects Into Tidy Data Frames. arXiv [stat.CO].

91. Wickham H. 2010 Stringr: Modern, consistent string processing. R J. 2, 38.

92. Xu S, Chen M, Feng T, Zhan L, Zhou L, Yu G. 2021 Use ggbreak to Effectively Utilize Plotting Space to Deal With Large Datasets and Outliers. Front. Genet. 12, 774846.

93. Lüdecke D, Ben-Shachar M, Patil I, Waggoner P, Makowski D. 2021 Performance: An R package for assessment, comparison and testing of statistical models. J. Open Source Softw. 6, 3139.

94. Lüdecke D. In press. sjPlot: Data visualization for statistics in social science. R package version

95. Long JA. In press. jtools: Analysis and presentation of social scientific data. R package version

96. Lenth R, Singmann H, Love J, Buerkner P, Herve M. 2019 Package ‘emmeans’.

97. Hothorn T, Bretz F, Hothorn MT. In press. The multcomp package. The R Project for Statistical Computing

98. Graves S, Piepho HP, Selzer L, Dorai-Raj S. In press. multcompView: Visualizations of Paired Comparisons. R package version 0. 1-7. R Found. Stat. Comput., Vienna, Austria

99. Wickham H, Wickham MH, RColorBrewer I. 2016 Package ‘scales’. R package version 1.

100. Schloerke B, Crowley J, Cook D, Briatte F, Marbach M. In press. GGally: extension to ‘ggplot2’. R package version 1.4. 0. R Foundation for Statistical

101. Electronic supplementary information for Hensley et al. 2023. Collective synchrony of mating signals modulated by ecological cues and social signals in bioluminescent sea fireflies. https://github.com/NikoHensley/ESM_egd_collective_behavior

