## Supplemental Results, Figures, and Tables for "Collective synchrony of mating signals modulated by ecological cues and social signals in bioluminescent sea fireflies"

### Supplementary Information

### Results

#### Collective responses to intensity *ex situ*

In addition to stimulus duration, EGD groups also show sensitivity in their collective response to stimulus intensity (Fig. S6, Table S3), responding readily to low levels of light. We observe that a relatively low level of light (estimated ~1,319.91 W sr^-1 m^-2 s, see Methods) elicits a maximum group response, and above which there is little increase in the number of responding males. We lack absolute measurements for the brightness of conspecific displays, and therefore cannot compare the intensity of our stimulus to that of ecologically-relevant social stimuli. At least in wavelength, our *ad hoc* stimulus mimics the colour of conspecific displays, with a peak emission frequency (lambda max) at ~450 nm (Fig. S5), and which is similar to bioluminescent emission spectra from crushed EGD at ~468 nm [[39]](https://paperpile.com/c/wrgnty/mtM0), as well as the visual sensitivity of congeneric luxorine ostracods at ~473 nm [[101]](https://paperpile.com/c/wrgnty/oQcq).

### Discussion

#### On variation in the number of pulses per display

Although males differ in the number of pulses in each display train (Fig. S3), we caution against an energetic interpretation of this variation, as we currently have no metric of the physiological demands during signalling nor do we know if there are fitness consequences for variation in the number of pulses during mate choice. Previous work indicates that the energetics of bioluminescent signalling in ostracods may be a relatively small amount of total luminescent ability [[102]](https://paperpile.com/c/wrgnty/0bEl), but this study could not account for the metabolic activity of swimming. From an information theory perspective, an increased number of pulses coincides with a larger ‘duty cycle’ or relatively more ‘on’ time, although species differ in the mean and maximum number of pulses per display (Fig. 1B, Table 1; [[37,103]](https://paperpile.com/c/wrgnty/s8ru+stYu)). Anecdotally, individuals within the same species found across habitats that vary in depth produce signals of varying pulse number despite high within species stereotypy, suggesting that multiple ecological factors can limit signal form in ostracods like EGD.

### Methods

#### Measuring stimulus intensity

Because we co-opted an a*d hoc* stimulus, intensity was not measurable in standardised units during experiments, and may or may not accurately model conspecific or environmental stimuli. Instead, we used a relative percentage of power output to regulate the intensity of the stimulus; by using an Arduino, we could pass up to a maximum voltage output (2 V), which could also be modulated in a standardised manner relative to this maximum. We used stimuli from ~2% to 100% of the electric potential across 10 experiments, and which in Arduino code are expressed in bytes: 5 (1.9% or 0.038 V), 10 (3.9% or 0.078 V), 20 (7.8% or 0.156 V), 30 (11.8% or 0.236 V), 50 (19.6% or 0.392 V), 70 (27.5% or 0.55 V), 90 (35.3% or 0.706 V), 120 (47.1% or 0.942 V), 180 (70.6% or 1.412 V), 255 (100% or 2 V).

Post-hoc measurements indicated that our stimuli were maximally: 966 to 30,117 W·sr^−1^·m^−2^ for stimuli at 5 to 255 bytes of power, respectively, and measured directly in front of the stimulus (Fig. S5). We measured side-welling spectral irradiance of the LED covered with the blue plastic bag using a fibre-optic spectrometer based on an Ocean Optics USB2000 (Dunedin, FL, USA). We calculated intensity by measuring the irradiance of the stimulus from 300 - 800 nm at different power levels (5, 25, 50, 100, 200, 255 bytes) at three locations in the aquaria, using a 1000 um fibre fitted with a cosine corrector (CC-3) and calibrated with a NIST (National Institute of Standards and Technology) traceable tungsten halogen lamp (LS-1, Ocean Optics), as done previously [[104]](https://paperpile.com/c/wrgnty/FC69).

References

101.	Huvard AL. 1993 Analysis of visual pigment absorbance and luminescence emission spectra in marine ostracodes (Crustacea: Ostracoda). *Comp. Biochem. Physiol. A Physiol.* **104**, 333–338.

102.	Rivers TJ, Morin JG. 2012 The relative cost of using luminescence for sex and defense: light budgets in cypridinid ostracods. *J. Exp. Biol.* **215**, 2860–2868.

103.	Cohen AC, Morin JG. 1993 The cypridinid copulatory limb and a new genus Kornickeria (Ostracoda: Myodocopida) with four new species of bioluminescent ostracods from the Caribbean. *Zool. J. Linn. Soc.* **108**, 23–84.

104.	Escobar-Camacho D, Pierotti MER, Ferenc V, Sharpe DMT, Ramos E, Martins C, Carleton KL. 2019 Variable vision in variable environments: the visual system of an invasive cichlid (Cichla monoculus) in Lake Gatun, Panama. *J. Exp. Biol.* **222**. (doi:[10.1242/jeb.188300](http://dx.doi.org/10.1242/jeb.188300))

105.	Moen RA, Pastor J, Cohen Y. 1999 Antler growth and extinction of Irish elk. *Evol. Ecol. Res.* **1**, 235–249.

##### Table S1. Linear model results predicting the time between collective events by the # of days after the full moon, the year of the field season, the # of minutes after sunset, and their interaction.

|  | **Time between collective events (s)** | | | | |
| --- | --- | --- | --- | --- | --- |
| *Predictors* | *Estimates* | *CI* | *Statistic* | *p* | *df* |
| (Intercept) | -338.67 | -482.17 – -195.17 | -4.65 | **<0.001** | 224 |
| # days after full moon | 5.3 | 1.36 – 9.24 | 2.65 | **0.009** | 224 |
| Year [2022] | 151.81 | -5.58 – 309.20 | 1.9 | 0.059 | 224 |
| # minutes post sunset | 9.53 | 6.82 – 12.25 | 6.92 | **<0.001** | 224 |
| Year x # min. post sunset | -4.33 | -7.25 – -1.41 | -2.92 | **0.004** | 224 |
| Observations | 229 | | | | |
| R^2^ / R^2^ adjusted | 0.438 / 0.428 | | | | |

##### Table S2. Results of a linear model describing changes in collective response to variation in stimulus duration (first and second order), tank used (a proxy for group-level effects), and the order in which tanks are tested.

|  | **# of displays** | | |
| --- | --- | --- | --- |
| *Predictors* | *Estimates* | *CI* | *p* |
| (Intercept) | 8.86 | 6.48 – 11.23 | **<0.001** |
| Stimulus duration [1st degree] | 49.90 | 40.44 – 59.35 | **<0.001** |
| Stimulus duration [2nd degree] | -58.35 | -67.80 – -48.90 | **<0.001** |
| Tank [B] | 7.61 | 3.07 – 12.15 | **0.001** |
| Tank [C] | 20.28 | 13.85 – 26.71 | **<0.001** |
| Tank [D] | 16.27 | 7.32 – 25.21 | **<0.001** |
| Tank [A] × Testing order | 1.57 | 0.75 – 2.39 | **<0.001** |
| Tank [B] × Testing order | -1.31 | -3.64 – 1.01 | 0.267 |
| Tank [C] × Testing order | -5.76 | -8.04 – -3.48 | **<0.001** |
| Tank [D] × Testing order | -3.80 | -6.24 – -1.36 | **0.002** |
| Observations | 257 | | |
| R^2^ / R^2^ adjusted | 0.563 / 0.547 | | |

##### Table S3. Results of a linear model describing changes in collective response to variation in stimulus intensity, tank used (a proxy for group-level effects), and the order in which tanks are tested.

|  | **# of displays** | | |
| --- | --- | --- | --- |
| *Predictors* | *Estimates* | *CI* | *p* |
| (Intercept) | 1.03 | -3.09 – 5.16 | 0.622 |
| radiance [log10] | 4.58 | 3.40 – 5.76 | **<0.001** |
| Tank [B] | 3.14 | -1.75 – 8.04 | 0.207 |
| Tank [C] | 10.86 | 3.46 – 18.26 | **0.004** |
| Tank [D] | 3.70 | 1.21 – 6.19 | **0.004** |
| Trial | -1.11 | -1.97 – -0.25 | **0.012** |
| Tank [A] × Testing order | 0.97 | 0.01 – 1.92 | **0.047** |
| Tank [B] × Testing order | 2.32 | -0.50 – 5.15 | 0.106 |
| Tank [C] × Testing order | -1.68 | -4.46 – 1.10 | 0.235 |
| Observations | 238 | | |
| R^2^ / R^2^ adjusted | 0.337 / 0.314 | | |

##### Table S4. Results of a linear model describing changes in individual latency to signal during stimulus presentation with respect to variation in stimulus duration, tank testing order, and distance from stimulus.

|  | **Latency to respond (s)** | | | |
| --- | --- | --- | --- | --- |
| *Predictors* | *Estimates* | *CI* | *p* | *df* |
| (Intercept) | 3.37 | 2.42 – 4.32 | **<0.001** | 193.00 |
| Stimulus duration (s) | 0.28 | 0.20 – 0.35 | **<0.001** | 193.00 |
| Testing order | 0.45 | 0.22 – 0.68 | **<0.001** | 193.00 |
| Distance from stimulus (cm) | -0.11 | -0.13 – -0.09 | **<0.001** | 193.00 |
| Observations | 197 | | | |
| R^2^ / R^2^ adjusted | 0.569 / 0.563 | | | |

##### Table S5. Results of a linear model describing changes in individual latency to signal after a stimulus presentation with respect to stimulus duration (first and second order), and the order in which tanks are tested.

|  | **Latency to respond (s)** | | | |
| --- | --- | --- | --- | --- |
| *Predictors* | *Estimates* | *CI* | *p* | *df* |
| (Intercept) | 0.80 | 0.67 – 0.93 | **<0.001** | 524.00 |
| Stimulus duration [1st degree] | -1.39 | -2.35 – -0.42 | **0.005** | 524.00 |
| Stimulus duration [2nd degree] | 3.98 | 3.01 – 4.95 | **<0.001** | 524.00 |
| Testing order | 0.06 | 0.01 – 0.12 | **0.033** | 524.00 |
| Observations | 528 | | | |
| R^2^ / R^2^ adjusted | 0.121 / 0.116 | | | |

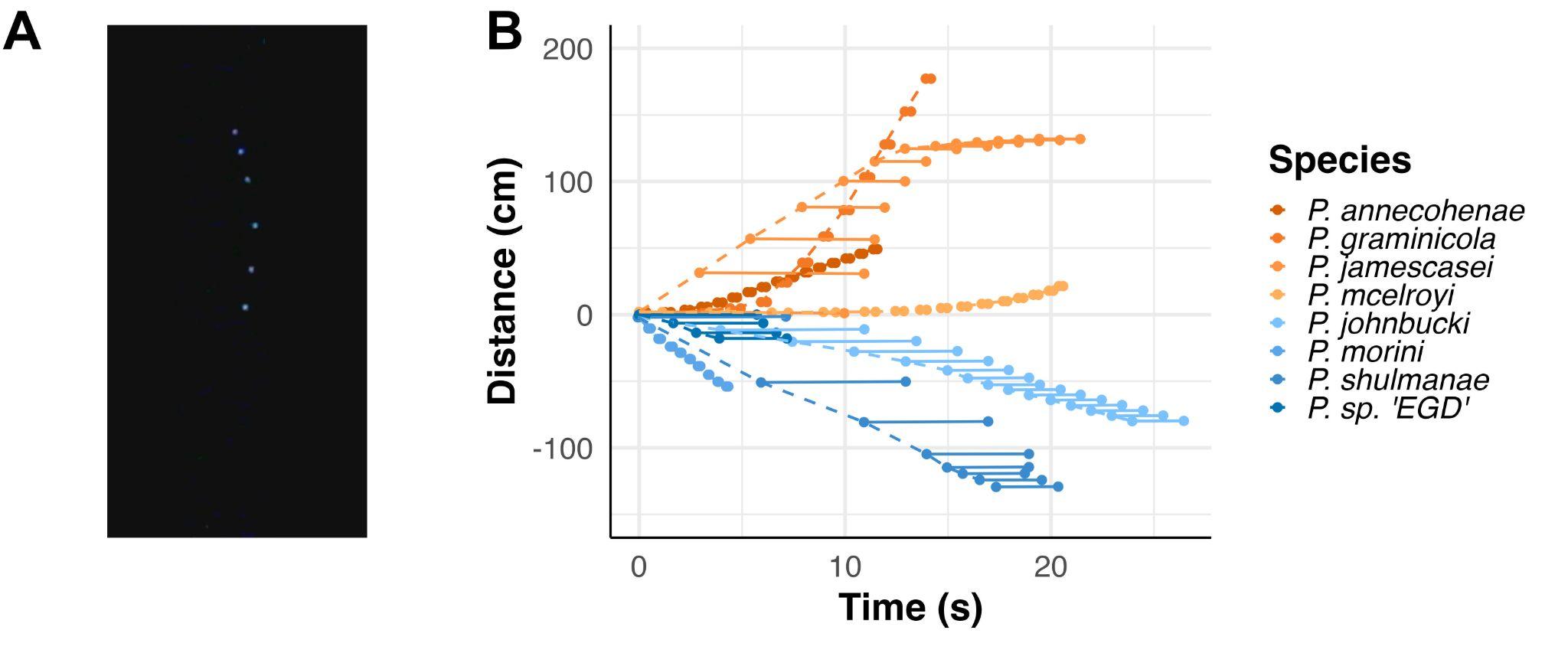

##### Figure S1. Time-distance plots of the average display train characteristics for *Photeros* species. Displays are coloured by species (note EGD in dark blue), and separated by whether pulses propagate downwards (blues) or upwards (oranges) over time within a display. Solid, horizontal lines connect the beginning and end of single pulses. Dashed lines connect the start of different pulses within the same display. Other species data from [[37]](https://paperpile.com/c/wrgnty/s8ru).

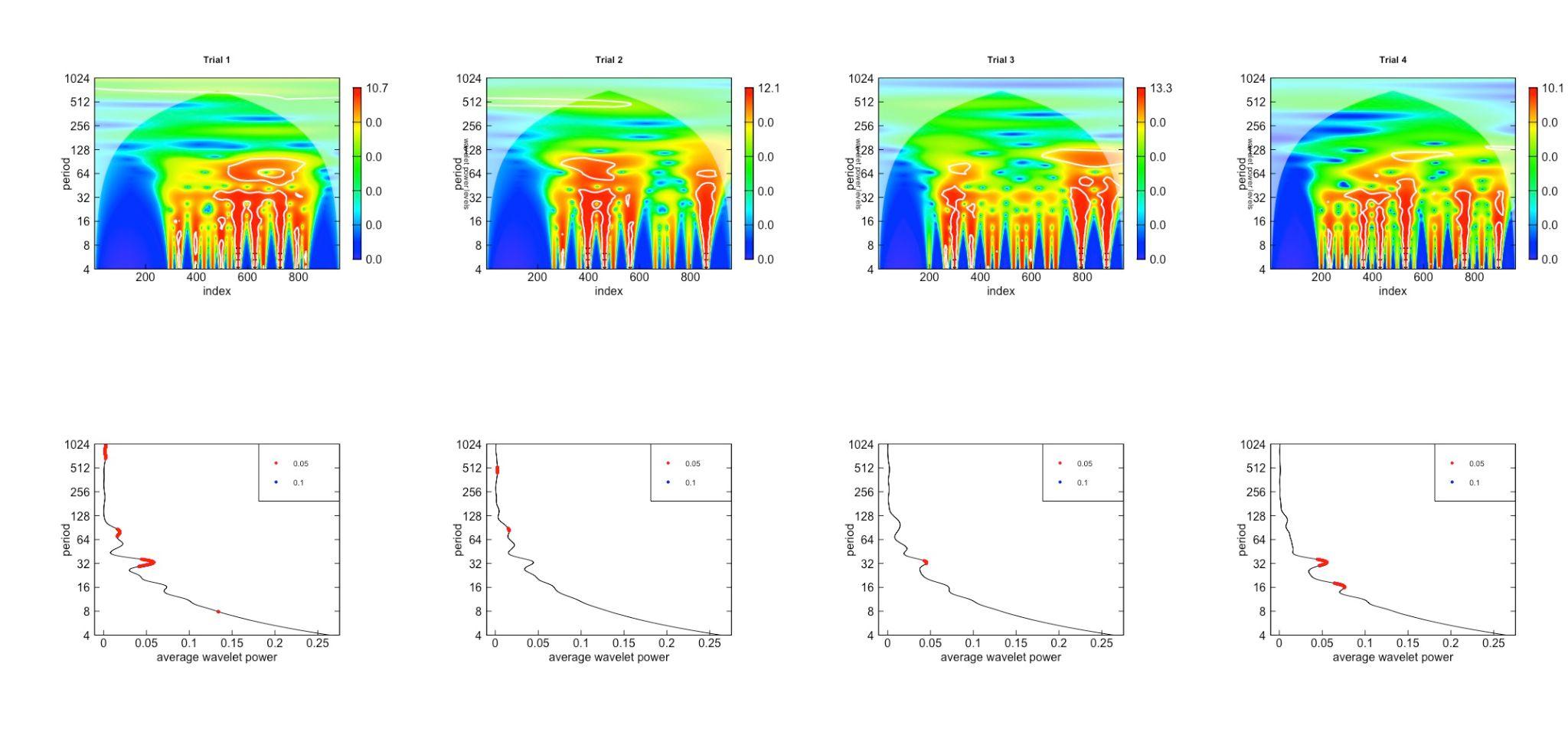

##### Figure S2. Top row: Wavelet transformation of count observations over time from Trials 1 - 4 of Fig. 4, from left to right respectively. Bottom row: Average wavelet power for each observation trial. Strong signals are found in lower periods for each, with a consistent periodicity every 32 sec, and with secondary periods at 16 and 64 seconds. Periods are measured in seconds, corresponding to the sampling interval of the time series.

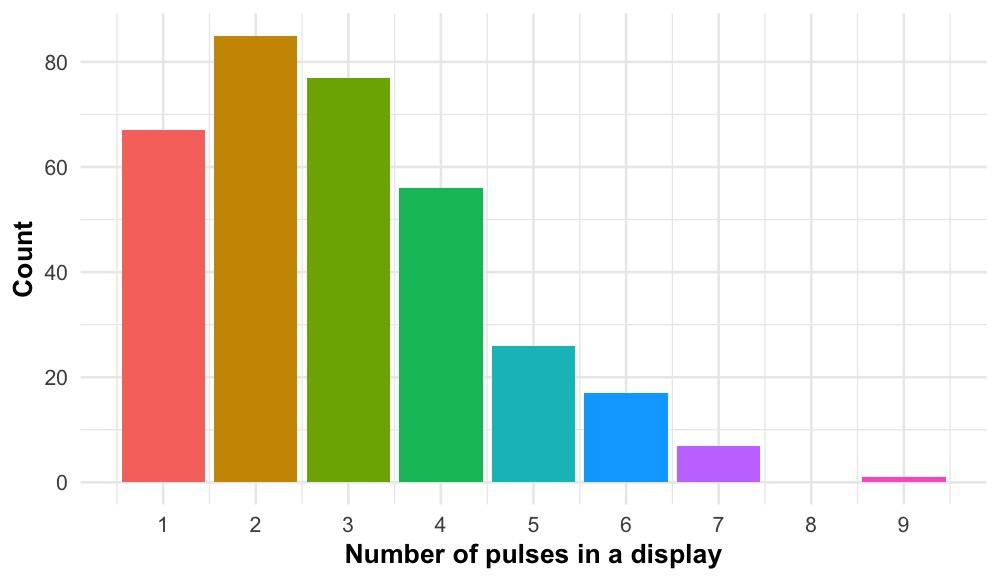

##### Figure S3. Histogram of variation in pulse number during *ex situ* observations (Trials 1 - 4). Signals with fewer pulses per display are more common, given the animal density (n = 100) and water depth (~20 cm) in the tanks.

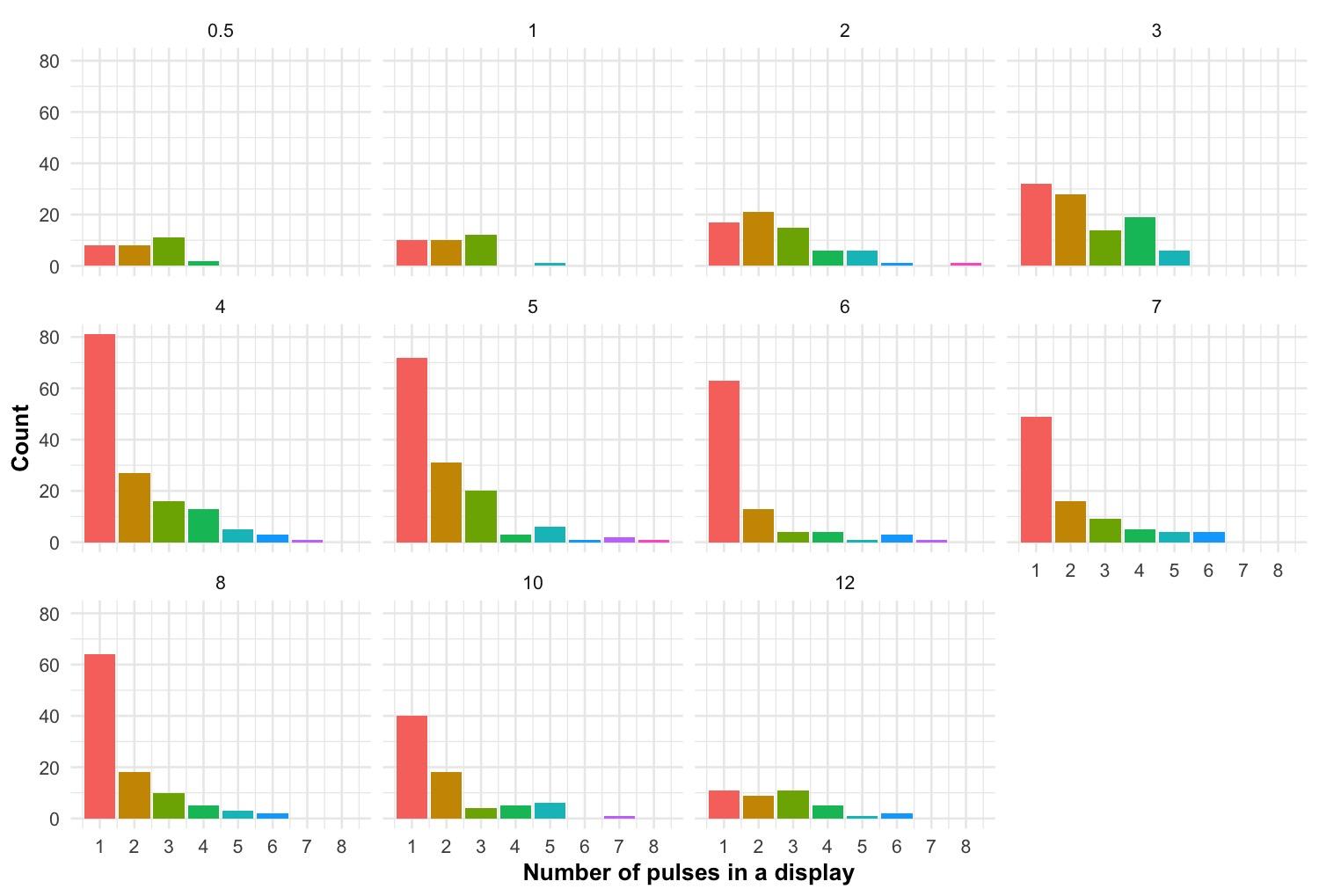

##### Figure S4. Change in the number of signals and distribution of pulse number per display as a function of the duration of the artificial stimulus. As we only have two annotated trials per stimulus, we cannot perform statistics on these data. We present here to corroborate our observations (Fig. 5A) that intermediate stimulus durations produce more signals.

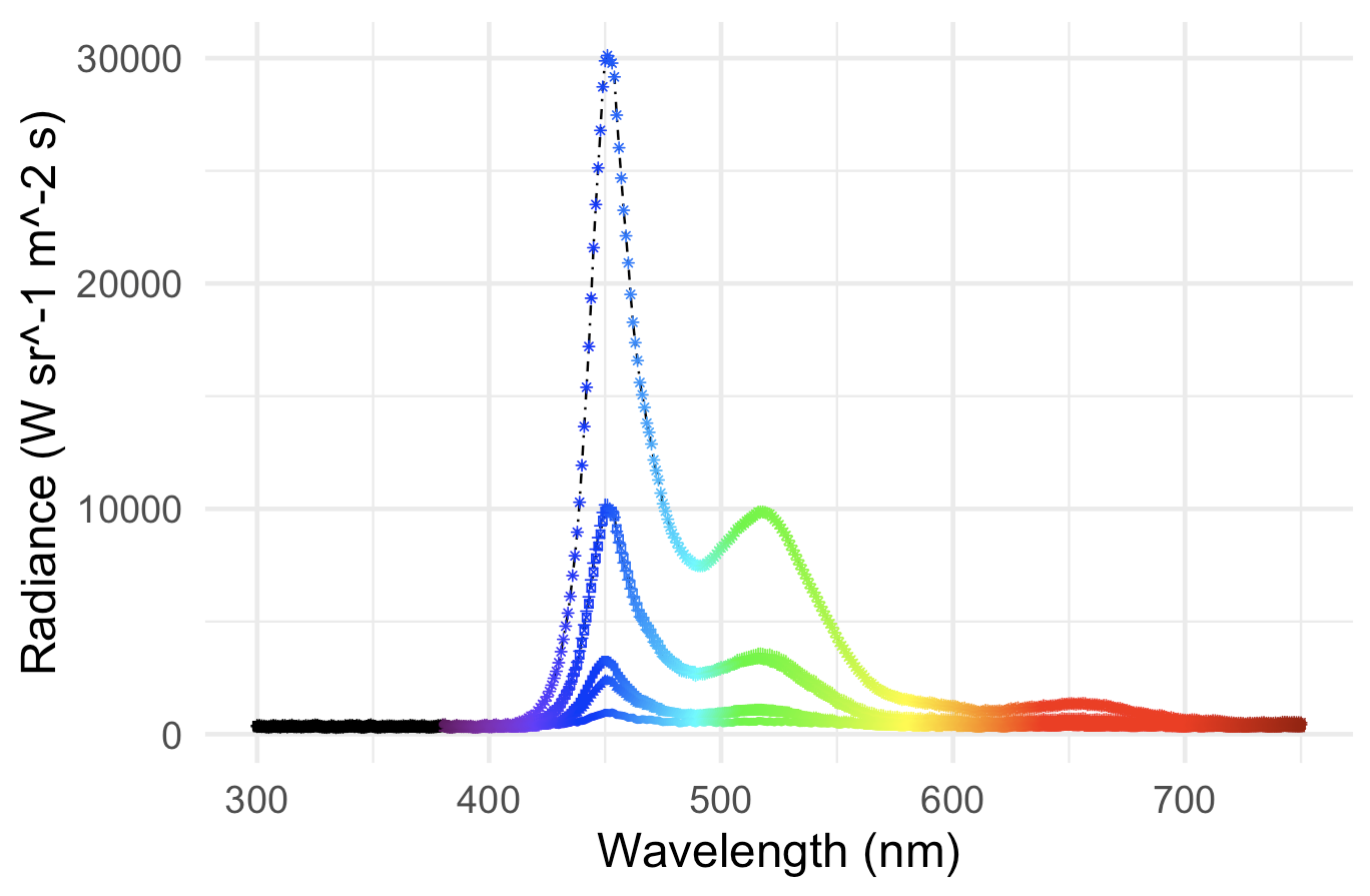

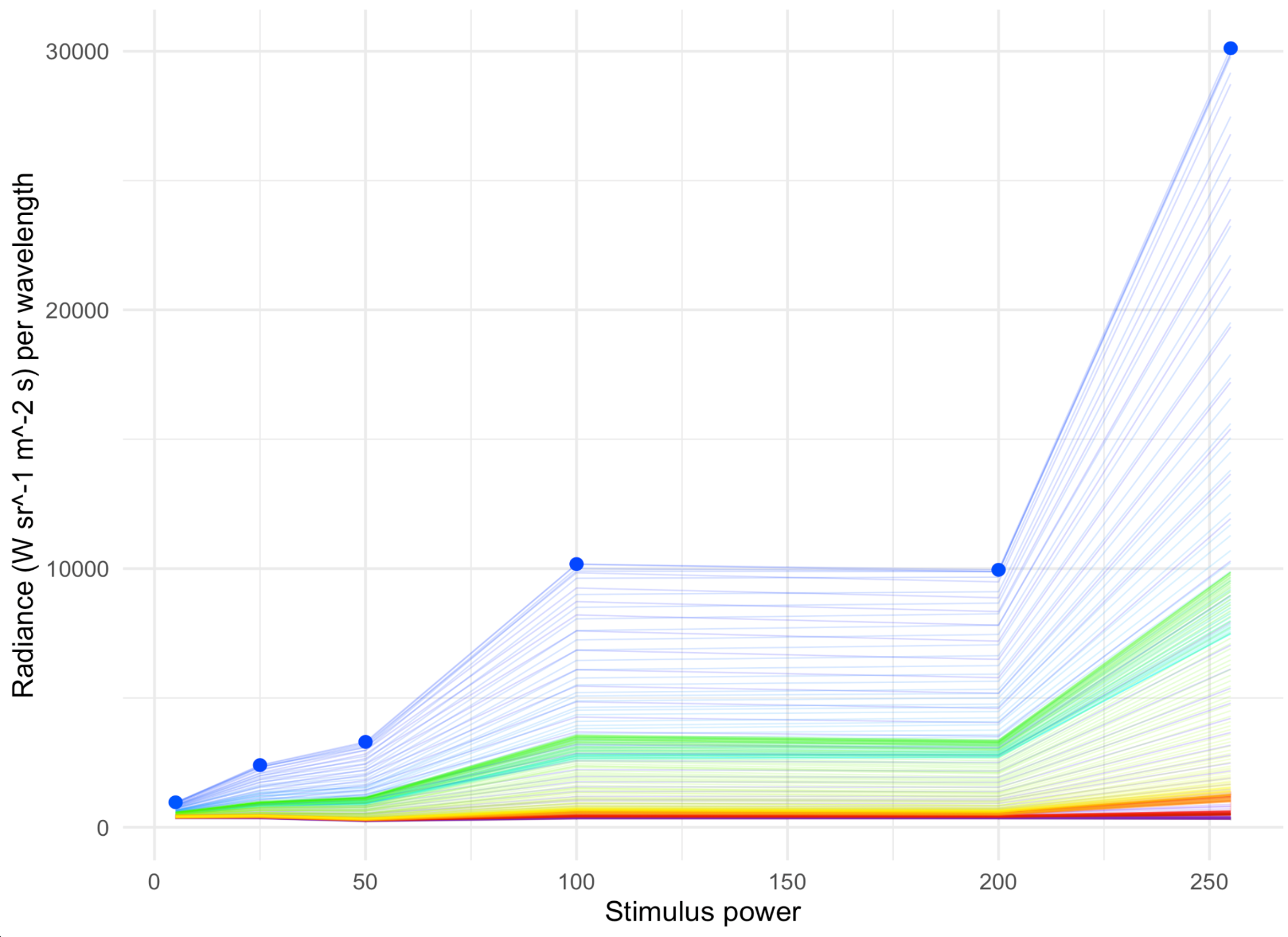

##### Figure S5. Distribution of radiance intensities across wavelengths for different power levels. Above: radiance by wavelength, with each line corresponding to a different power level. Below: same data as above, but plotted as radiance by power level for every wavelength measured. Therefore, each horizontal line in the plot below represents a vertical slice from the plot above.

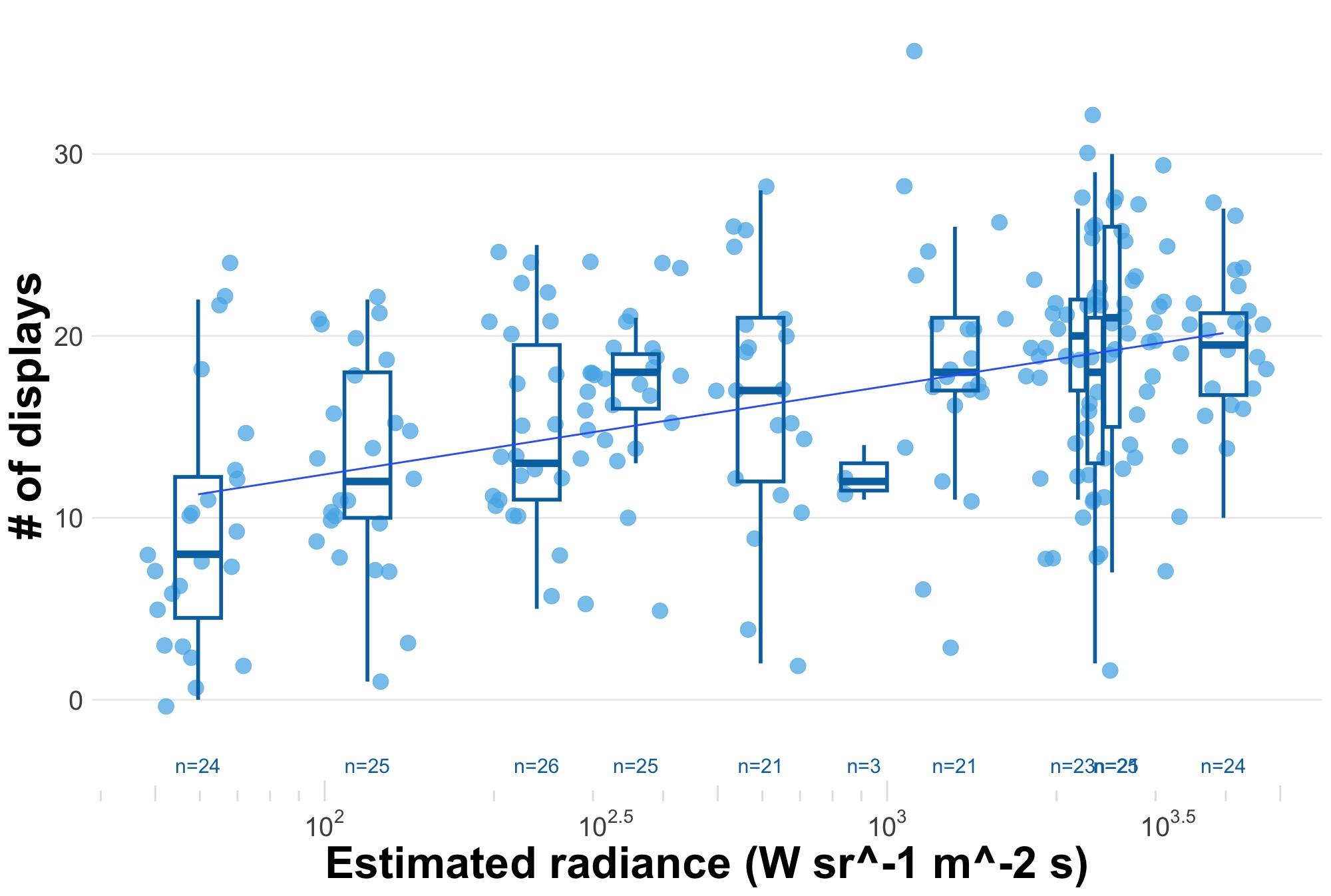

##### Figure S6. Groups of signalling males show variation in the magnitude of their collective responses to stimuli varying in intensity as measured in radiance. Semi-log plot demonstrates a plateau in response to changes in the stimulus radiance (estimated from data, see Methods and Fig. S3). Data are jittered for visualisation. Each datum represents a single experimental trial where we count by eye the number of individual displays produced immediately after stimulus onset. Regression is demonstrative; for model fit, see Table S2.

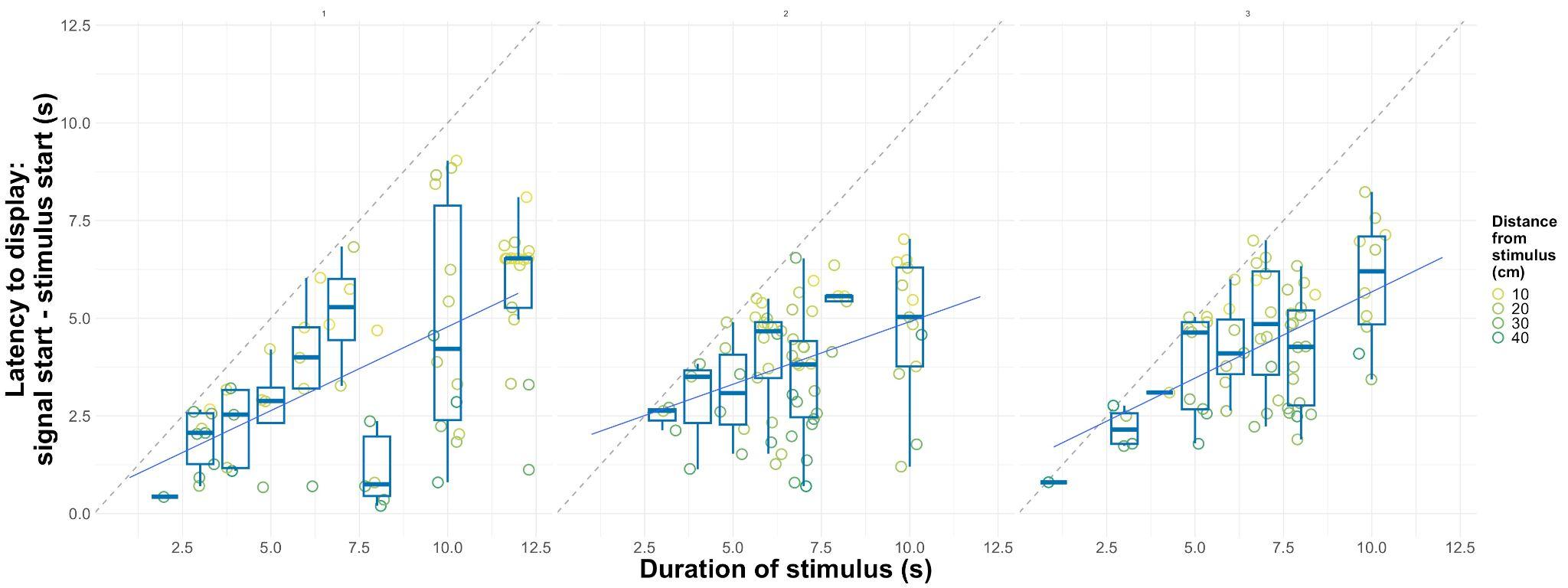

##### Figure S7. Latency to respond for males creating signals during a stimulus presentation as a function of distance stimulus duration, distance from stimulus, and the testing order of the tank during the night (left to right, panels 1 - 3). Grey dashed lines represent 1:1 line, which we would expect if males simply waited as long as possible until they responded. Instead, we see a regression with lower slope. Regression lines are demonstrative only; for the full model, see Table S3.

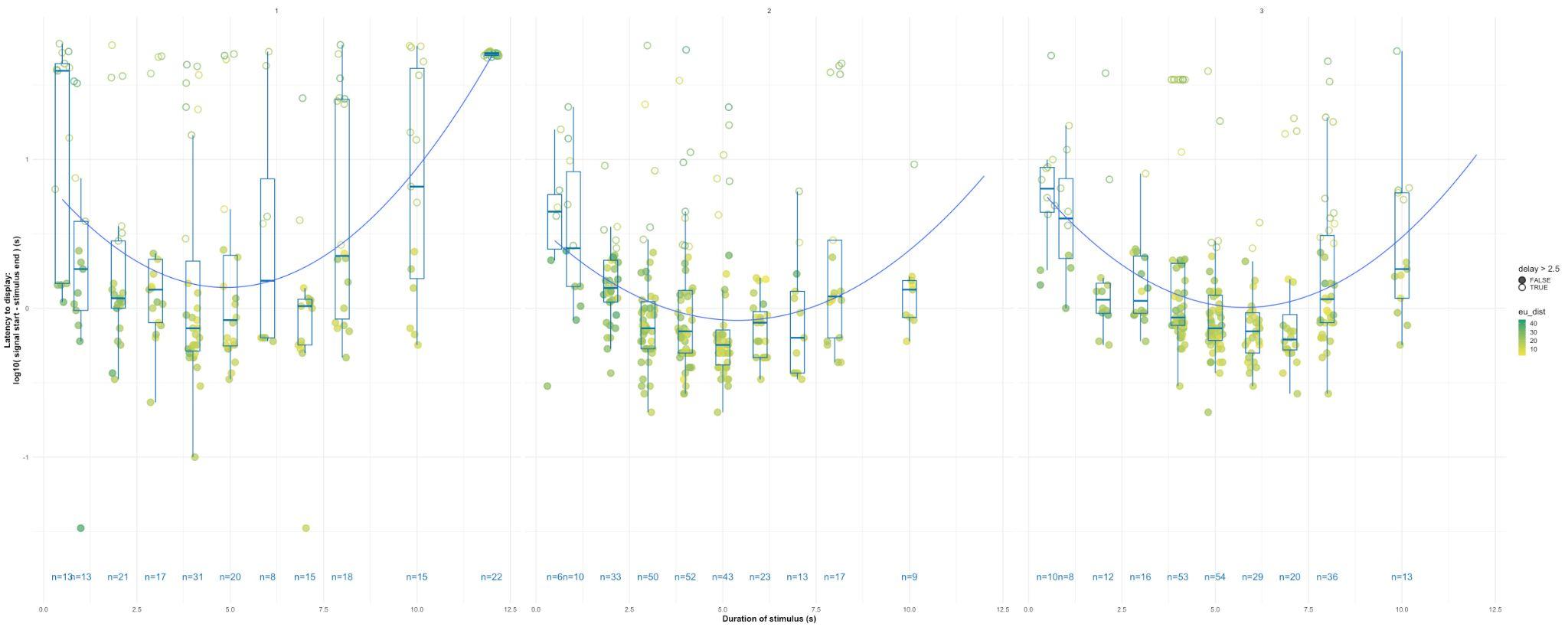

##### Figure S8. Latency to respond for males creating signals directly after a stimulus presentation ends across variation in stimulus presentation, all data. Note the log transformed y-axis. Pattern is similar to Fig. 5D, which is a dataset restricted to males responding within 2.5 s only (filled data here).

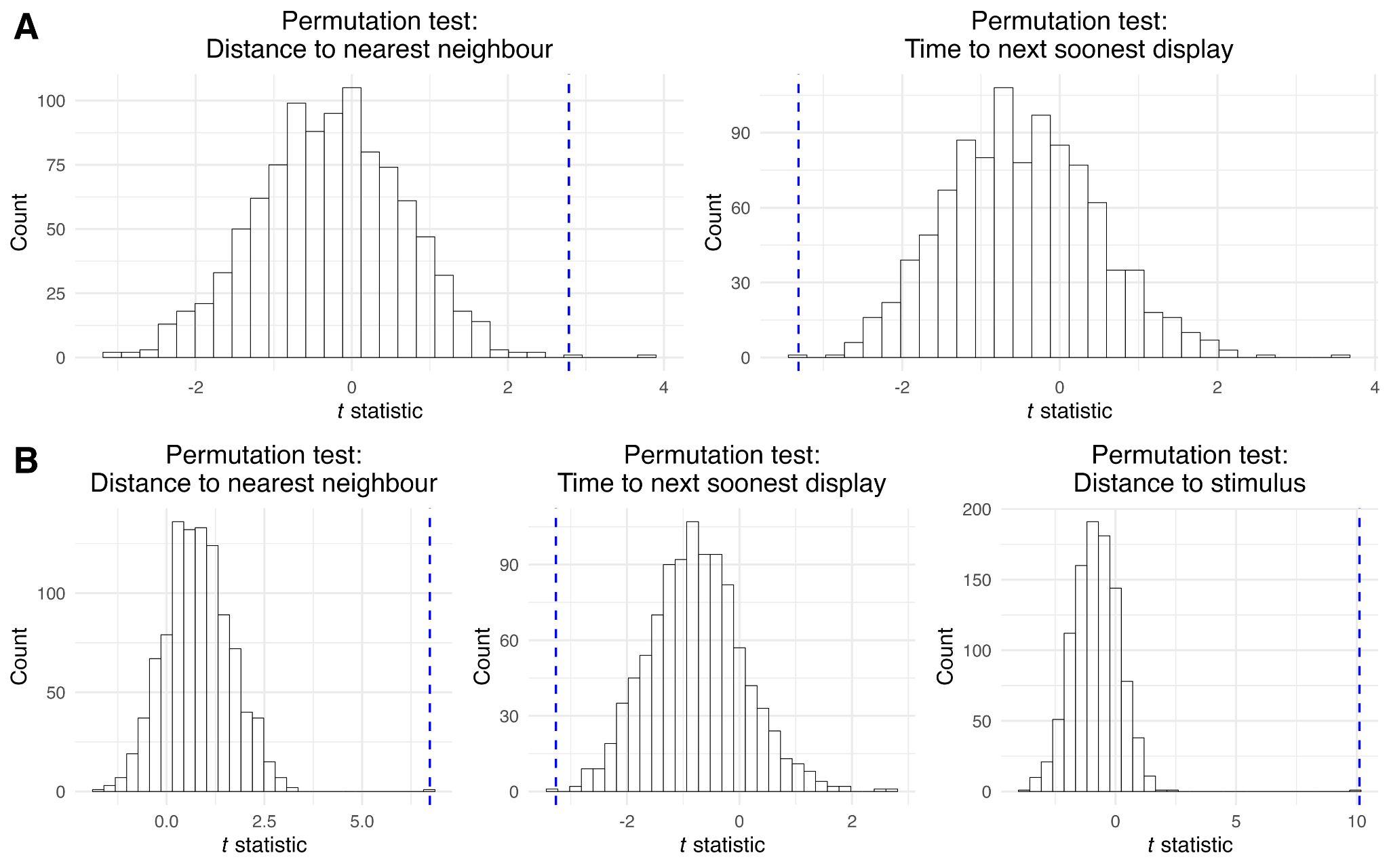

##### Figure S9. Results from the permutation tests for (A) linear models on observational trials, and (B) linear models on experimental trials. *t* statistic distributions are the result of 1000 linear models per predictor variable on the number of displays with different permutations of the datasets. Blue dashed lines represent the observed value for each predictor variable.

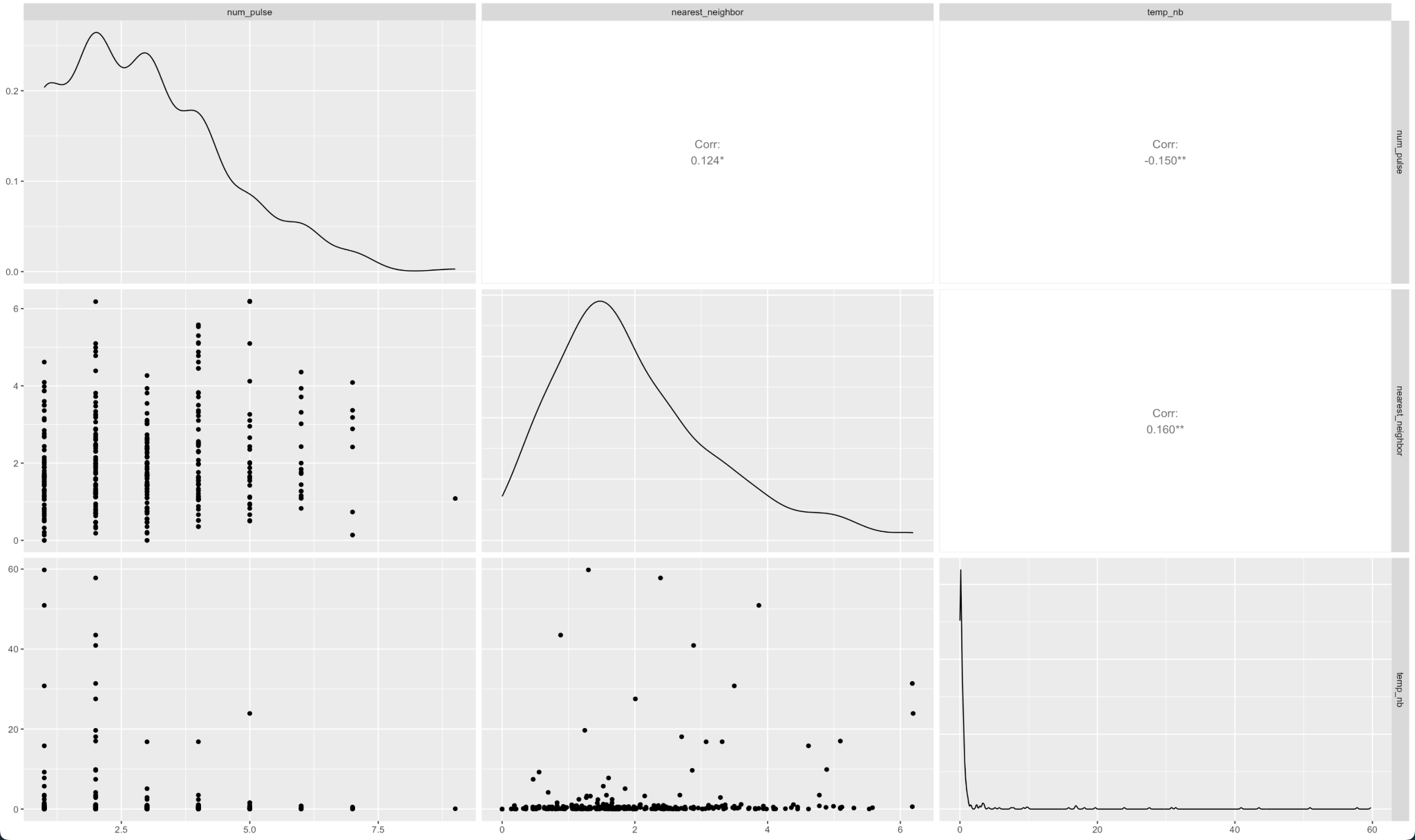

##### Figure S10. Correlations between time and distance variables in observational trials.

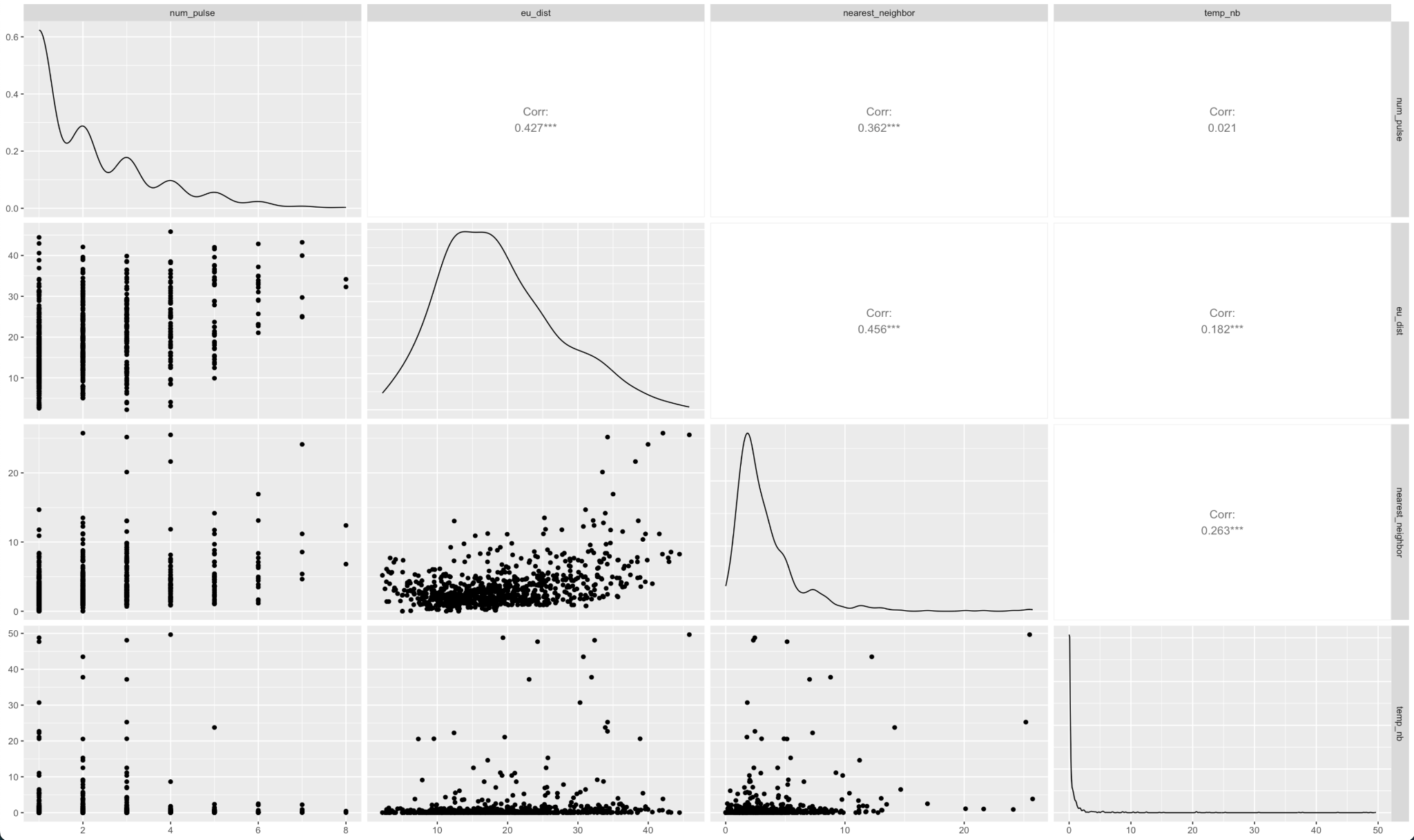

##### Figure S11. Correlations between time and distance variables in experimental trials.
